# In vivo anti-FAP CAR T therapy reduces fibrosis and restores liver homeostasis in metabolic dysfunction-associated steatohepatitis

**DOI:** 10.1101/2025.02.25.640143

**Authors:** Chittampalli N. Yashaswini, Bruno Cogliati, Tianyue Qin, Tran To, Thomas Williamson, Tyler E. Papp, Kenneth Li, Raisa Rasul, Li Chen, Adi Lightstone, Haig Aghajanian, Hamideh Parhiz, Shuang Wang, Joel G. Rurik, Jonathan A. Epstein, Scott L. Friedman

**Affiliations:** Division of Liver Diseases, Icahn School of Medicine at Mount Sinai; New York, NY, USA; Tisch Cancer Institute, Icahn School of Medicine at Mount Sinai; New York, NY, USA; Department of Oncological Sciences, Icahn School of Medicine at Mount Sinai; New York, NY, USA; The Precision Immunology Institute, Icahn School of Medicine at Mount Sinai; New York, NY, USA; Medical Scientist Training Program, Icahn School of Medicine at Mount Sinai; New York, NY, USA; Department of Pathology, School of Veterinary Medicine and Animal Science, University of São Paulo; São Paulo, Brazil; Department of Pharmacological Sciences, Icahn School of Medicine at Mount Sinai; New York, NY, USA; Department of Cell and Developmental Biology, Perelman School of Medicine at the University of Pennsylvania; Philadelphia, PA, USA; Penn Cardiovascular Institute, Perelman School of Medicine at the University of Pennsylvania; Philadelphia, PA, USA; Institute for Regenerative Medicine, Perelman School of Medicine at the University of Pennsylvania; Philadelphia, PA, USA; Department of Medicine, Perelman School of Medicine at the University of Pennsylvania; Philadelphia, PA, USA; Pharmanest, Inc; Princeton, NJ; Capstan Therapeutics, Inc; San Diego, CA; Center for Infectious Medicine, Department of Medicine Huddinge, Karolinska Institutet; Stockholm, Sweden

## Abstract

In this study, we aimed to determine the efficacy of in vivo chimeric antigen receptor (CAR) T cell therapy, generated by targeted lipid nanoparticles (t-LNPs), as an anti-fibrotic in metabolic dysfunction-associated steatotic liver disease. Hepatic fibrosis is a key predictor of mortality in liver disease, driven by fibrogenic hepatic stellate cells (HSCs). In heart, chimeric antigen receptor (CAR) T cells targeting fibroblast activation protein alpha (FAP) reduce murine cardiac fibrosis. However, the value of this approach in liver is unknown. We explored the anti-fibrotic potential of in vivo-generated anti-FAP CAR T cells in metabolic dysfunction-associated steatohepatitis (MASH), a highly prevalent disease with no approved anti-fibrotic therapies. We first established that FAP expression in both human and murine MASH is specific to HSCs. We then used flow cytometry, Sirius Red morphometry, digital pathology analysis, and single nuclear RNA-sequencing to assess the impact of anti-FAP CAR T cell therapy on murine MASH. Anti-CD5 targeted-LNPs carrying anti-FAPCAR mRNA generate activated, transient anti-FAP CAR T cells, which significantly reduced fibrosis by depleting pro-fibrogenic HSCs, and by modulating immune cells, endothelial cells and hepatocytes in a non-cell autonomous manner to mitigate inflammation and restore hepatic homeostasis. These findings reinforce the potential of in vivo CAR T therapy to attenuate a highly morbid and pervasive liver disease through integrated, multicellular salutary effects.

**One Sentence Summary:** RNA-based treatment transiently reprograms immune cells to target scar-forming cells in fatty liver disease, thus improving liver health overall.

## INTRODUCTION

Metabolic dysfunction-associated steatotic liver disease (MASLD), formerly known as non-alcoholic fatty liver disease (NAFLD), is the most prevalent chronic liver disease globally, affecting about one-third of adults(1, 2). The surge in MASLD prevalence mirrors the escalating rates of obesity, type 2 diabetes, and metabolic syndrome worldwide(3, 4). Metabolic dysfunction-associated steatohepatitis (MASH) is a serious outcome of MASLD developing in ∼20% of patients, and is characterized by inflammation, hepatocyte ballooning, and fibrosis, with a significant risk of progression to cirrhosis and hepatocellular carcinoma(5, 6).

Hepatic fibrosis, defined as an aberrant accumulation of extracellular matrix in the liver, leads to increased tissue stiffness, architectural distortion, impaired oxygenation, and compromised liver function(7, 8). Fibrosis is the leading prognostic indicator of patient mortality(9) in MASH, underscoring the urgent need for effective anti-fibrotic therapies.

To tackle fibrosis, clearing hepatic stellate cells (HSCs), the liver’s fibrogenic cell through their activation into myofibroblasts, has emerged as a potential strategy(10). Despite constituting only ∼5% of resident hepatic cells in healthy liver and 10-15% in diseased liver, HSCs wield significant influence over liver homeostasis and disease progression(11, 12). HSCs are quiescent within uninjured liver, where they store vitamin A and are essential to maintaining liver homeostasis through secretion of paracrine mediators that sustain hepatocyte health, proliferation and differentiation. In response to liver injury, HSCs transdifferentiate from a quiescent to an activated state, driving fibrogenesis through their enhanced proliferation, contractility, and extracellular matrix (ECM) production, among other cellular responses(13, 14).

With the advent of single cell technologies, it has become clear that HSCs have considerable heterogeneity beyond the two stages of quiescence and activation(15). Therefore, strategies to attenuate effects of activated HSCs necessitate precise targeting strategies to mitigate fibrosis without compromising normal liver function(16-19). As an example, global depletion of HSCs prevents liver regeneration through loss of HSC-derived mitogenic signals to hepatocytes(20). Thus, selective depletion of only disease-causing HSC subsets is essential(10, 21). Selective elimination of fibrogenic HSCs may provide more significant disease resolution than current therapies which target a single mediator or receptor, which remain clinically underwhelming.

A promising cell surface protein for HSC targeting is fibroblast activating protein alpha (FAP). FAP is a serine protease of the dipeptidyl peptidase (DPP) family, which is primarily expressed on the surface of activated fibroblasts, including HSCs in fibrotic microenvironments(22, 23). It plays a crucial role in modulating extracellular matrix (ECM) remodeling in fibrosis by cleaving ECM components, including collagen(24), and contributes to the activation and perpetuation of fibrogenic signaling pathways in HSCs through paracrine interactions with macrophages(25), thereby facilitating fibrosis progression. Thus, targeting FAP directly could enhance ECM remodeling and also alter the immunoregulatory status of macrophages within fibrotic tissues. Furthermore, FAP protease activity results in cleavage of fibroblast growth factor 21 (FGF21), a hormone primarily secreted by liver and adipose tissues which has salutary effects on the liver, including reduction of hepatic steatosis, inflammation, lipotoxicity and ER stress(26). For this reason, FGF21 analogues and agonists are in development for use in diseases like MASH(27). Thus, FAP targeting in patients could amplify FGF21 signaling and improve hepatocyte health.

In cancer, selective cell depletion using chimeric antigen receptor (CAR) T cell therapy is an established, cutting-edge immunotherapeutic that has garnered significant success(28, 29) and is now gaining traction in non-oncological diseases(10, 30-32). However, for the latter indications, a persistent CAR T cell population as observed in conventional ex vivo engineered CAR T cells(33), is undesirable. To address this concern and improve patient accessibility, CAR T cells can be generated in vivo by delivering the mRNA instructions for CAR generation within targeted-lipid nanoparticles (t-LNPs) targeting T lymphocytes. By leveraging mRNA, this transient approach avoids integration into the T cell’s genome and therefore unwanted persistence(30). This strategy was originally developed to generate anti-FAP CAR T cells to reverse fibrosis in a mouse model of heart disease(30), but its relevance to other fibrotic diseases is uncertain.

In this study we aim to determine the impact of in vivo anti-FAP CAR T cells on hepatic fibrosis in a well-validated mouse model of MASH, determining not only their effect on fibrogenic cell targets but also the collateral impact on resident liver cells and infiltrating immune cells.

Here we show that using T cell-directed t-LNPs to deliver CAR-encoding anti-FAP mRNA leads to depletion of profibrotic HSCs with reduction of hepatic fibrosis. Using a multi-modal approach including flow cytometry, immunostaining, immunoblotting, digital pathology analysis, and single-nucleus RNA sequencing (snRNAseq), we demonstrate successful anti-FAP CAR T generation in vivo, degranulation of CD8^+^ anti-FAP CAR Ts in the liver, and a striking reduction in fibrosis and pro-fibrogenic HSC subsets. HSC depletion exerts non-cell autonomous effects on other hepatic cells, including a significant increase in activated Ly6c^lo^ monocyte-derived macrophages, which promote liver fibrosis resolution, a preservation of healthy phenotype of liver sinusoidal endothelial cells (LSECs), and downregulation of aberrant lipid regulation in hepatocytes, including reduced hepatocyte expression of cytochrome P450 omega-hydroxylase (*Cyp4a14*), which contributes to progression of MASLD(34). Altogether, these data highlight the exciting potential of t-LNP-based CAR T therapy to treat hepatic fibrosis and reverse MASLD pathogenesis.

## RESULTS

### FAP expression is restricted to fibrogenic HSCs and increased in MASH

To determine if FAP is a suitable target for in vivo CAR T therapy in liver, we evaluated its expression in murine and human MASH. In the mouse Fibrosis and Tumor (FAT) MASH model, which recapitulates human MASH(35) (Fig 1A), FAP and glial fibrillary acidic protein (GFAP), a marker of activated HSCs in the liver, were co-localized (Fig 1B). FAP staining increased in murine MASH liver at progressive timepoints compared to healthy chow-fed livers, with a significant increase in early, 6-week FAT MASH (Fig 1C-D). Similar analyses of human liver confirmed elevated FAP expression in MASH tissue compared to healthy liver (Fig S1A, B). Analysis of published snRNAseq dataset of the 24-week timepoint of the FAT MASH model(36) confirmed *Fap* expression by activated HSCs (HSC2), as well as by some quiescent HSCs (HSC1) (Fig 1E, G). Sub-clustering of the HSCs from this dataset(37) reinforces the significant heterogeneity of HSCs, pinpointing a small subcluster arising between the quiescent and activated states, termed “quiescent precursors of activated HSCs” (Fig 1F and (37)). In mice, *Fap* is most highly expressed by activated HSCs and quiescent precursors of activated HSCs subsets (Fig 1F, H). While the dendritic cell (DC) cluster in snRNAseq data of murine FAT MASH appeared to highly express *Fap* mRNA, this expression is over-represented due to limited capture of immune cells by snRNAseq.

**Fig. 1.**
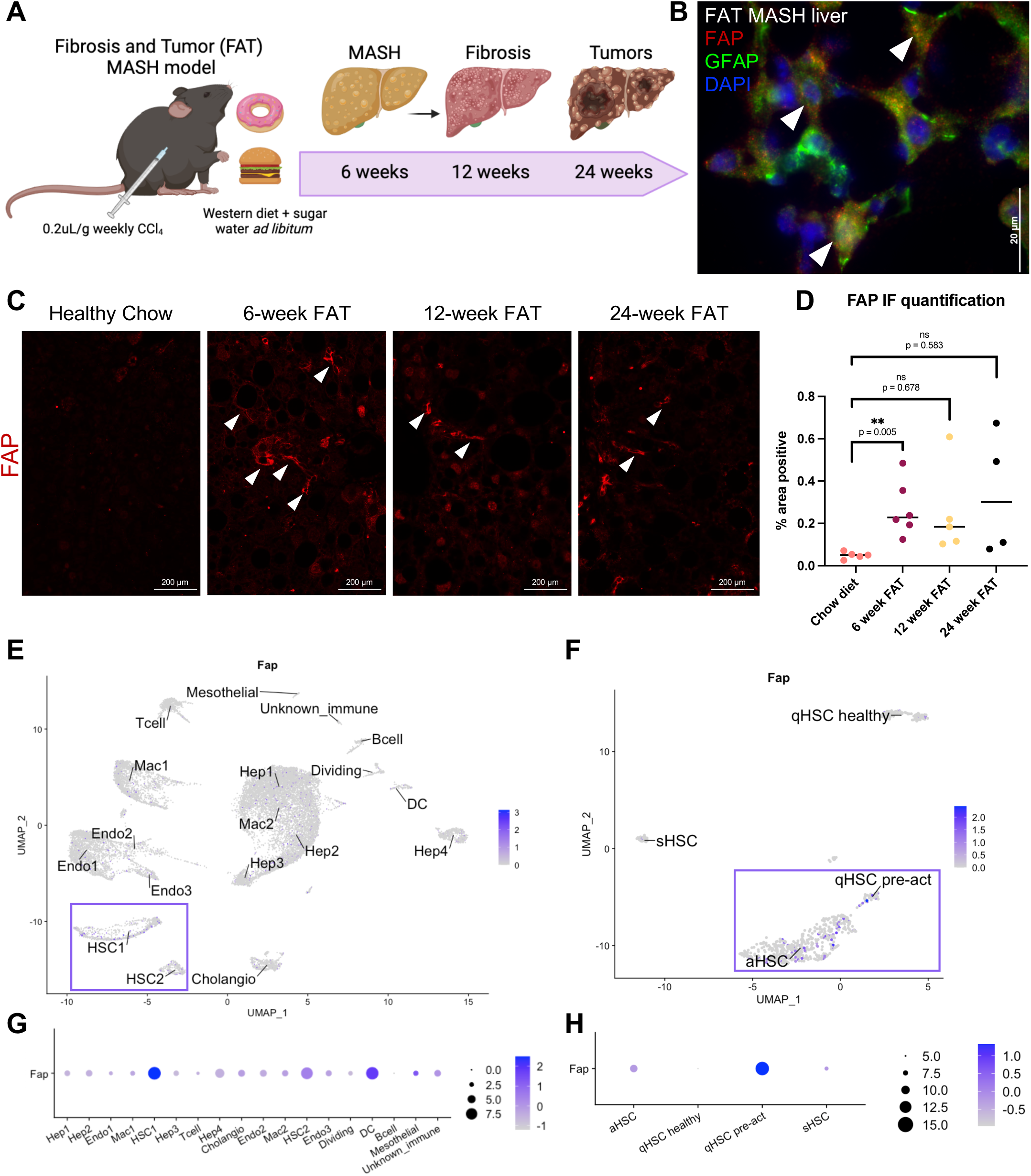
FAP is expressed by activated HSCs in murine MASH. **(A)** Fibrosis and Tumor (FAT) murine MASH model setup. **(B)** Co-immunostaining of 6-week FAT liver with FAP (red), GFAP (green) and DAPI (blue), where white arrows indicate cells with co-staining. **(C)** FAP immunostaining of livers from chow vs FAT mice, where white arrows indicate areas of FAP positivity. **(D)** FAP fluorescence quantification. **(E)** snRNAseq Feature Plot of *Fap* expression in FAT liver. **(F)** snRNAseq Feature Plot of Fap expression in FAT HSC subsets. **(G)** Dot Plot of FAP expression in FAT liver. **(H)** Dot Plot of FAP expression in FAT HSC subsets.

Analysis of snRNAseq datasets of patient MASH versus healthy liver tissue(36) highlights the specificity of *FAP* to HSCs in humans (Fig S1C, E), and a subsequent analysis of subclustered human HSCs(37) indicates that *FAP* is primarily expressed by activated HSCs in MASH patients (Fig S1D, F). These data establish the specificity of FAP to fibrogenic HSC subsets in murine and human MASH. Furthermore, the distinct elevation of FAP expression in human MASH tissues, and its specificity to activated HSCs reaffirms its strong potential as a therapeutic target to selectively deplete fibrogenic HSCs, while sparing quiescent, non-fibrogenic HSCs.

### Targeted-LNPs are specific for CD5^+^ cells and generate functional CAR T cells

Mice were injected with targeted-LNPs (t-LNPs) or vehicle at the 6-week FAT MASH timepoint, aligned with the time at which there was significantly elevated FAP expression (Fig 1C) and accumulating fibrosis(36). A group of age-matched chow-fed mice injected with vehicle were also included as an additional control. This therapeutic treatment model was chosen instead of a prophylactic model, to mirror potential treatment strategy in established human MASH. Mice were injected with CD5-directed t-LNPs carrying mCherry mRNA (CD5-mCherry), CD5-directed t-LNPs carrying anti-FAP chimeric antigen receptor (CD5-FAPCAR) mRNA, or a water vehicle (Fig 2A). Alternatively, in a similar, separate experiment, mice were injected with IgG-coated (non-targeting) t-LNPs carrying FAPCAR mRNA (IgG-FAPCAR), CD5-FAPCAR t-LNPs, or a saline vehicle, to validate CD5 as an LNP target, and provide added validation of the overall results (Fig S2A). Eighteen hours after t-LNP administration, extracellular CD5 was significantly reduced in the livers and spleens of CD5-FAPCAR recipients compared to IgG-FAPCAR recipients (Fig S2B, C), indicating that anti-CD5 LNPs successfully engaged cell surface CD5 and were endocytosed (gating strategy in Fig S7A). The trending reduction in extracellular CD5 in the blood leukocytes (Fig S2B, C), is also consistent with liver accumulation of FAPCAR^+^ cells by 18 hours post-LNP injection.

**Fig. 2.**
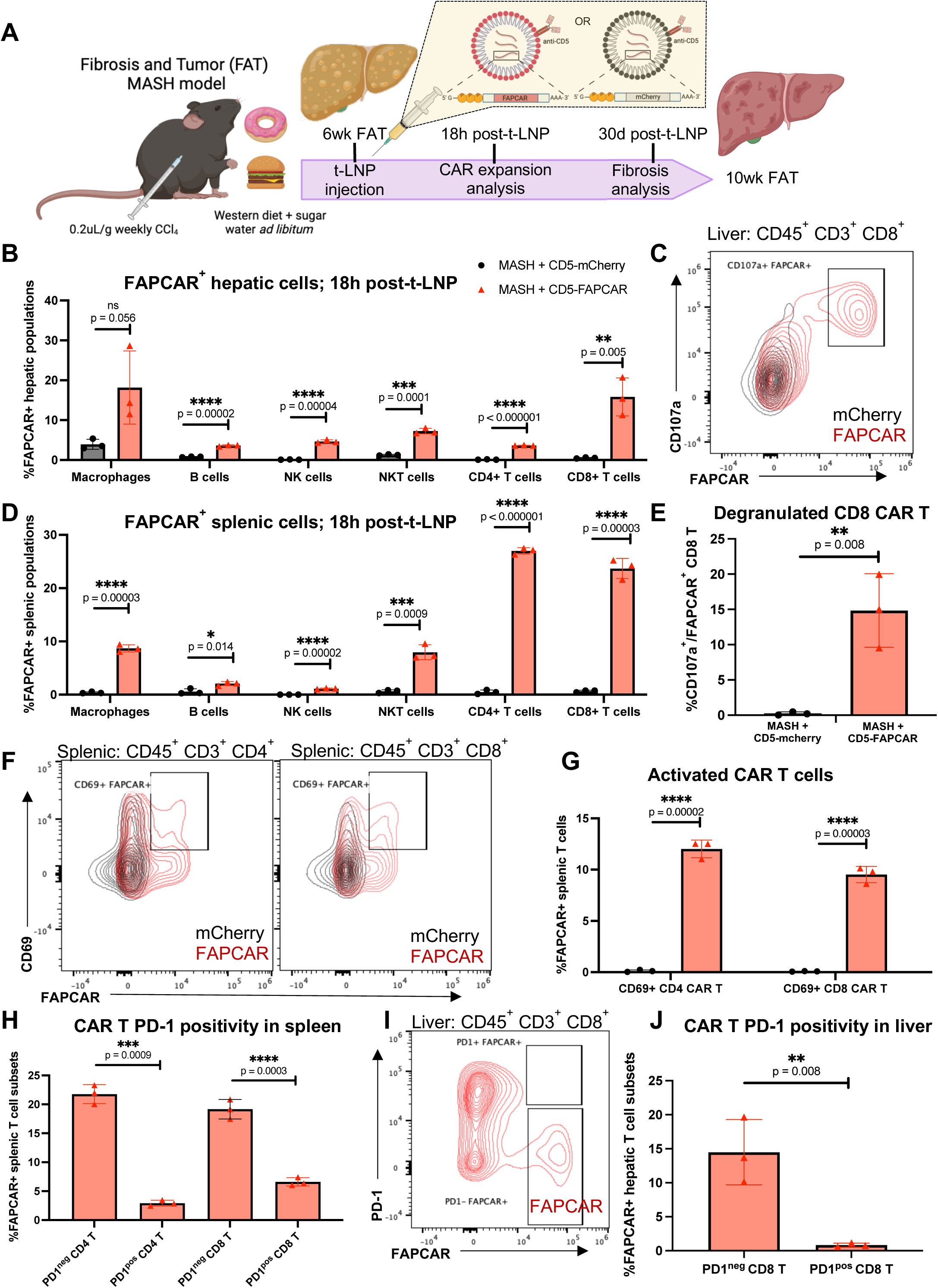
in vivo anti-FAP CAR^+^ cells include non-exhausted CD8 T cells that show evidence of effector function. **(A)** Experimental setup, where treatment is CD5-FAPCAR t-LNP and control is CD5-mCherry t-LNP. **(B)** Flow cytometry quantification of FAPCAR^+^ hepatic immune subsets. **(C)** Flow plot of hepatic CD8 CAR T cells expressing extracellular CD107a 18 hours after t-LNP injection. **(D)** Flow cytometry quantification of FAPCAR^+^ splenic immune subsets 18 hours after t-LNP injection. **(E)** Quantification of hepatic CD8 CAR T cells expressing extracellular CD107a 18 hours after t-LNP injection, with CD5-mCherry in black and CD5-FAPCAR in red. **(F)** Flow plots and **(G)** quantification of splenic CD4 (left) and CD8 (right) CAR T cells expressing CD69 18 hours after t-LNP injection, with CD5-mCherry in black and CD5-FAPCAR in red. **(H)** Flow quantification of PD-1^neg^ vs PD-1^pos^ splenic CD4 (left) and CD8 (right) CAR T cells in CD5-FAPCAR mice 18 hours after t-LNP injection. **(I)** Flow plot and **(J)** quantification of PD-1 positivity in FAPCAR^+^ CD8 T cells of CD5-FAPCAR recipients 18 hours after injection. In all bar graphs, CD5-mCherry results are graphed with black circles, and CD5-FAPCAR results are graphed with red triangles.

Eighteen hours after t-LNP injection, FAPCAR^+^ immune cells were detectable in livers (Fig 2B, Fig S2D) and spleens (Fig 2D, Fig S2E) of CD5-FAPCAR recipients, with a plurality of CAR T cells (gating strategy in Fig S7B). Approximately 5-25% of CD4 T cells and 15-25% of CD8 T cells were FAPCAR^+^. Immunostaining of splenic tissue demonstrated FAPCAR co-localization with CD8 and CD4 in the CD5-FAPCAR t-LNP recipients, with minimal co-localization in the controls, reinforcing the high specificity of the anti-CD5 LNPs to generate CAR T cells in vivo (Fig S2F). Hepatic CD8 FAPCAR^+^ T cells also stained positive for extracellular CD107a, also known as LAMP-1. When on the cell surface, CD107a is evidence of degranulation and indicates recent effector function, likely due to the presence of the CAR target - FAP - in injured liver (Fig 2C, E). Furthermore, approximately 10% of CD4 and CD8 CAR T cells expressed CD69, a marker that is rapidly induced on the cell surface after TCR/CD3 engagement, indicating successful activation of in vivo-generated CAR T cells (Fig 2F, G).

MASH livers are known to harbor exhausted T cells compared to healthy livers, which has posed challenges to immunotherapeutic approaches for primary cancers arising in this disease(38). However, in spleen and liver most CAR T cells are PD-1^neg^ (Fig 2H-J), indicating that in vivo CAR T cells are not exhausted, despite their generation in the context of MASH.

Altogether, anti-CD5 FAPCAR LNPs are highly specific CD5+ cells and generate in vivo anti-FAP CAR T cells in the spleen and liver that are non-exhausted, activated, and have effector function.

### In vivo anti-FAP CAR T therapy reduces liver fibrosis within 4 weeks

We observed a significant reduction in hepatic fibrosis in the CD5-FAPCAR recipients 30 days after t-LNP injection, as determined by Sirius Red staining (Fig 3A, B, Fig S3A, B), and immunofluorescence staining for aSMA, a marker of activated HSCs (Fig S3C, D). Digital pathology analysis of Sirius Red staining was performed by FibroNest^TM^, a high-resolution single-fiber AI-based platform that quantifies the histological phenotype of fibrosis severity. It classifies collagen fibers as “fine” or assembled” based on the complexity of the collagen networks, with assembled fibers being more intricately reticulated (containing >25 branches) and characterizes architectural complexity of fibrosis at a single collagen fiber level(39). This analysis revealed a reduction of assembled fibers and fibrosis architectural complexity in anti-CD5 FAPCAR t-LNP recipients (Fig 3C). The analysis also revealed a significant reduction in the *Phenotypic Fibrosis Composite Score*, which is an overarching phenotypic quantification of the progression of the fibrosis severity, and a significant reduction in the *Assembled Fibrosis Morphometric Composite Score*, which describes the collective morphometric traits of each individual assembled fiber, including fiber length, width, area, and perimeter (Fig 3D). By immunoblotting and blinded analysis, expression of COL1A1 and aSMA proteins in CD5-FAPCAR liver tissue were significantly reduced compared to CD5-mCherry liver tissue (Fig 3E-G), altogether indicating reduced fibrosis with in vivo anti-FAP CAR T therapy.

**Fig. 3.**
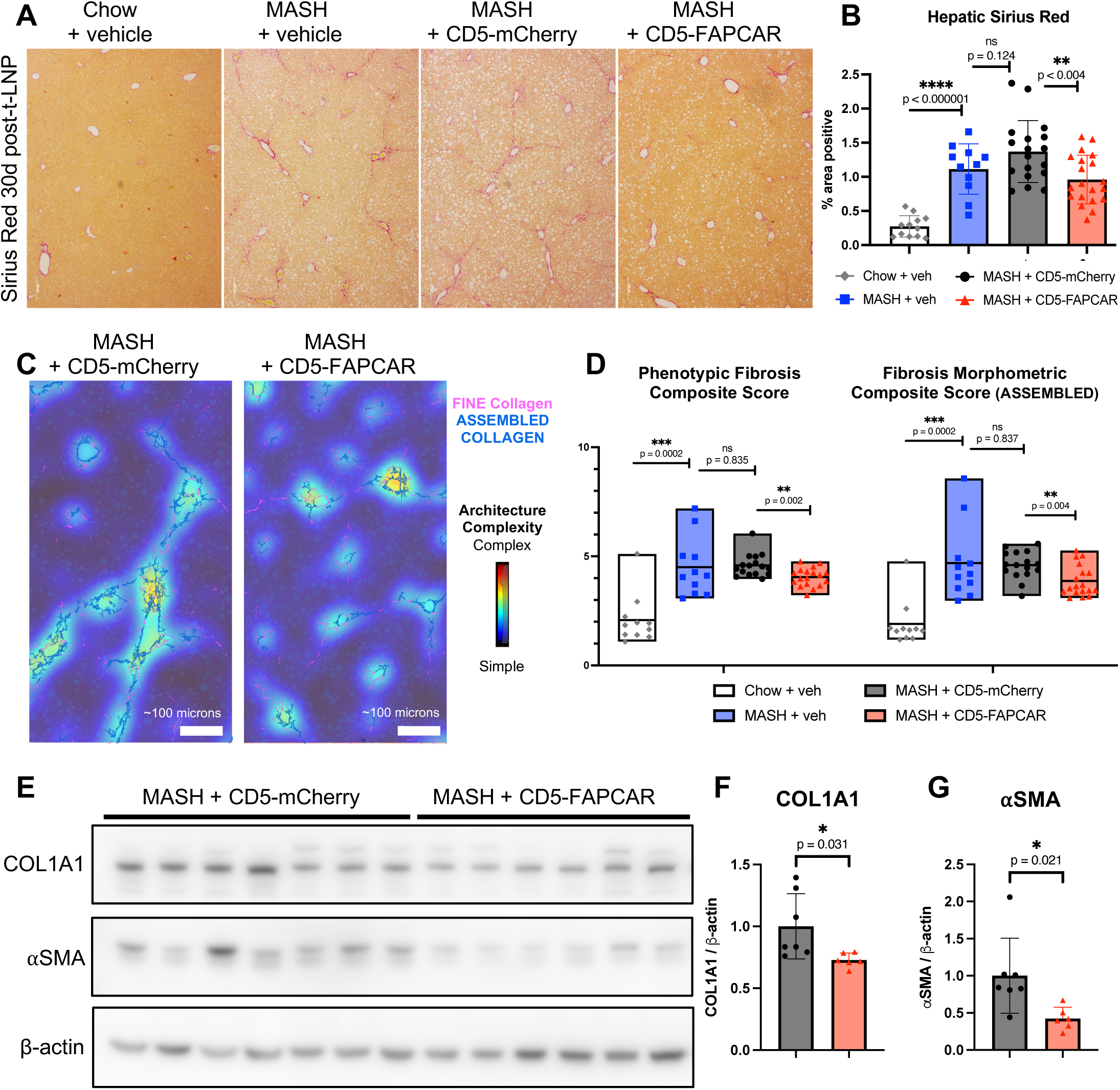
in vivo anti-FAP CAR T cells reduce hepatic fibrosis in MASH. **(A)** Sirius Red staining and **(B)** quantification 30 days after t-LNP injection. **(C)** Digital pathology analysis via FibroNest^TM^ platform of fine collagen (pink) and assembled collagen (blue), and fibrosis architectural complexity (color map) overlaid on Sirius Red. **(D)** Digital pathology quantification of fibrosis severity via FibroNest^TM^. **(E)** Immunoblotting and **(F, G)** quantification of COL1A1, ⍺SMA relative to β-actin.

### Anti-FAP CAR T cells target fibrogenic HSCs and redistribute HSC subsets

Flow cytometry analysis of CD45^-^ CD31^-^ aSMA^+^ nonparenchymal liver cells revealed a significant reduction in aSMA^+^ cells in CD5-FAPCAR recipients, compared to CD5-mCherry controls (Fig 4A, B) (gating strategy in Fig S7C). These data aligned with immunofluorescence data, which showed a trending reduction of aSMA (Fig 4C), altogether consistent with the depletion of fibrogenic HSCs following in vivo anti-FAP CAR T therapy.

**Fig. 4.**
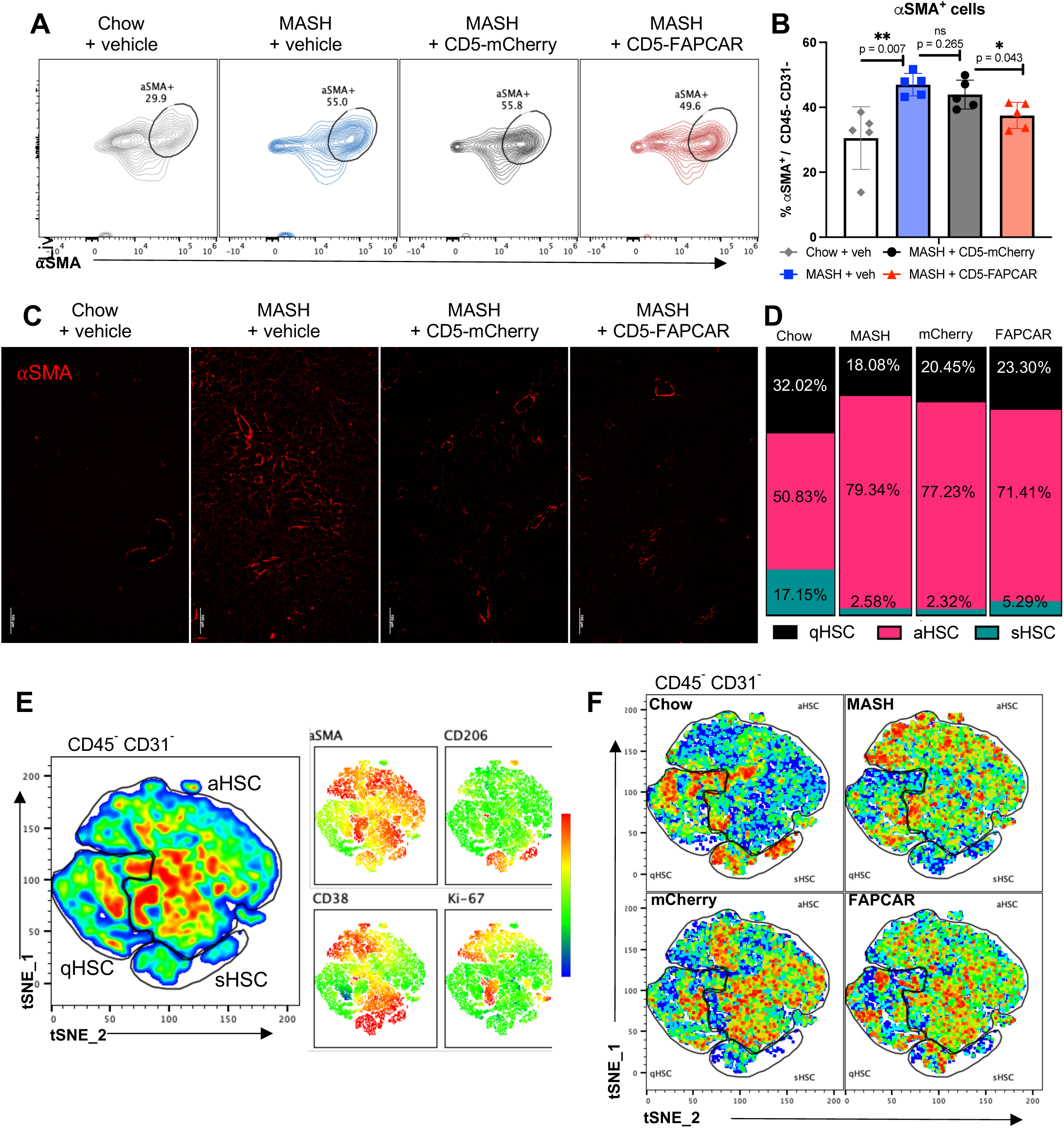
in vivo anti-FAP CAR T cells deplete fibrogenic HSCs, causing redistribution of HSC subsets. **(A)** Flow plots and **(B)** quantification of CD45^-^ CD31^-^ aSMA^+^ cells. **(C)** ⍺SMA staining. **(D)** Quantification of relative proportions of suggested HSC subsets from (F). **(E)** Flow tSNE plot of CD45^-^ CD31^-^ cells from all four experimental groups 30 days after injection (Chow + vehicle, MASH + vehicle, MASH + CD5-mCherry, MASH + CD5-FAPCAR), with suggested HSC subset gating and corresponding heatmaps showing distribution of selected markers. **(F)** Flow tSNE plots of each experimental group.

Flow cytometry tSNE analysis of HSC markers within the CD45^-^ CD31^-^ liver cell fraction allowed for identification of putative quiescent (qHSC), activated (aHSC) and senescent (sHSC) subsets (Fig 4E). Activated HSCs were determined by aSMA positivity; within the aHSCs; some were noted to be proliferative (Ki-67^+^), and/or expressed CD38 (a regulator of HSC activation, present in more severe fibrosis)(35, 40). Based on our recent study characterizing markers of HSC senescence in the FAT MASH model(37), cells expressing both aSMA and CD206 were classified as senescent. Remaining cells that did not express activation or proliferation markers were classified as quiescent. This tSNE analysis of HSC subsets demonstrated that healthy chow-fed mice treated with vehicle and MASH mice treated with CD5-FAPCAR had the largest proportion of qHSCs (32.02% and 23.03% respectively) and sHSCs (17.15% and 5.29% respectively), with the smallest proportion of aHSCs (50.83% and 71.41%, respectively). On the other hand, MASH mice that received vehicle or CD5-mCherry had the smallest proportion of qHSCs (18.08% and 20.45% respectively) and sHSCs (2.58% and 2.32% respectively), and the largest proportion of aHSCs (79.34% and 77.23% respectively) (Fig 4D, F). These data indicated that in vivo anti-FAP CAR T cells lead to a favorable redistribution of HSC subsets that reduces fibrogenic HSCs and increases anti-fibrotic HSCs.

snRNAseq clustering (Fig S4A, S8A-F) and analysis of livers 30 days post-t-LNP injection confirmed that *Fap* is specifically expressed by HSCs (Fig S4B), and that *Fap* expression is reduced in CD5-FAPCAR recipients compared to IgG-FAPCAR controls (Fig S4C). Further subclustering of HSCs based on previously published papers(17, 37) revealed distinct clusters (Fig S4D). Additionally, the relative frequencies of these HSC subclusters were altered, with reduced quiescent precursors of activated HSC and activated HSC subsets in CD5-FAPCAR recipients, and an increase in the quiescent and senescent HSC subsets compared to IgG-FAPCAR recipients (Fig S4E), aligning with flow cytometric data, and suggesting that in vivo anti-FAP CAR T therapy is efficacious at eliminating the pro-fibrogenic cells, while sparing non-fibrogenic quiescent HSCs, which are important for liver homeostasis.

### Anti-FAP CAR T therapy reprograms non-HSC hepatic cells, aiding MASH resolution

FAP expression by HSCs promotes HSC and macrophage profibrogenic activity in liver fibrosis(25). Flow cytometric analysis of immune populations 30 days after t-LNP injection showed a significant increase in activated (CD69^+^, CD86^+^) Ly6C^lo^ monocyte-derived macrophages in the liver in CD5-FAPCAR recipients compared to CD5-mCherry recipients (Fig 5A, B). While there were no significant changes in Ly6C^hi^ monocyte-derived macrophages, this observed shift in macrophage immunoregulatory status is likely beneficial, as Ly6C^lo^ monocyte-derived macrophages are “non-classical” and known to reduce hepatic inflammation and promote fibrosis resolution(41, 42). Additionally, Ly6C^lo^ monocyte-derived macrophages maintain the stability and health of the LSECs(43), which are typically damaged, dysfunctional and reduced in injured MASH liver(44). Flow cytometry analysis of CD45-CD31+ liver cells, presumed to be LSECs, reveals an increase in this population in CD5-FAPCAR recipients compared to CD5-mCherry recipients (Fig 5C). Comparison of the snRNAseq endothelial cell populations in the IgG-FAPCAR controls versus the CD5-FAPCAR treatments (Fig 5D) revealed five key differentially expressed genes (DEGs). Genes upregulated in CD5-FAPCAR recipients (*Atp5l* and *Glul*) are important for maintenance of vascular development and tone in endothelial cells(45, 46), whereas genes upregulated in the IgG-FAPCAR recipients (*Fmnl2*, *St6galnac3*, *Insr*) are implicated in loss of mechanical tissue homeostasis, pathologic angiogenesis, and cancer development(47-50) (Fig 5E, F). The expression pattern of these DEGs highlights that the CD5-FAPCAR endothelial cells are more similar to those of the healthy chow mice, whereas the IgG-FAPCAR endothelial cells are more similar to those of the diseased MASH mice. This suggests that recovery of the LSEC population and restoration of homeostatic endothelial cell function is one component of the observed MASH resolution following in vivo anti-FAP CAR T therapy.

**Fig. 5.**
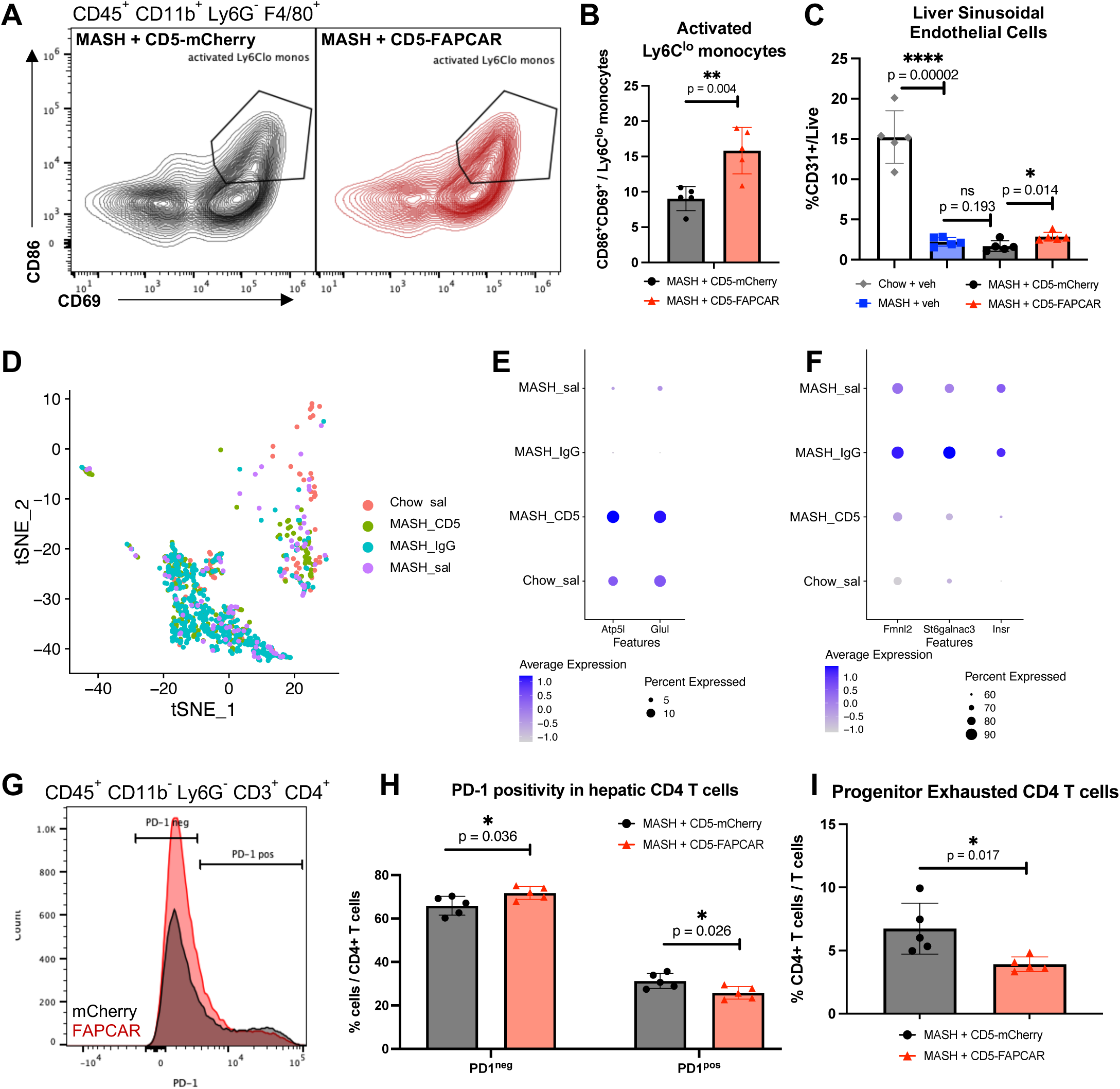
anti-FAP CAR T cells promote MASH resolution via modulation of hepatic immune cells and recovery of endothelial cells. **(A)** Flow plots and **(B)** quantification of CD45^+^ CD11b^+^ Ly6G^-^ F4/80^+^ Ly6C^lo^ that are CD86^+^ CD69+, detected 30 days after t-LNP injection. **(C)** Quantification of LSECs (CD45^-^ CD31^+^) cells detected 30 days after t-LNP injection. **(D)** snRNAseq tSNE plot of endothelial cells. **(E)** DotPlot of top differentially expressed genes that are elevated in CD5-FAPCAR. **(F)** DotPlot of top differentially expressed genes that are elevated in IgG-FAPCAR. **(G)** Flow plot of PD-1 positivity in the hepatic CD4 T cell fraction 30 days after t-LNP injection and **(H)** quantification. **(I)** Flow quantification of progenitor exhausted CD4 T cells (PD-1^low^, CD69^-^, Ki-67^+^).

Hepatic T cell exhaustion is elevated in MASH livers and implicated in the progression of disease(51). Flow cytometry analysis of PD-1 levels revealed a significant increase in PD-1^neg^ CD4 T cells, and a significant decrease in PD-1^pos^ CD4 T cells in CD5-FAPCAR recipients 30 days post-t-LNP injection (Fig 5G, H). Further analysis of exhausted CD4 T cell subsets identified a significant reduction in progenitor exhausted CD4 T cells, which are PD-1^low^, Ki-67^+^ and CD69^-^ (Fig 5I). This progenitor exhausted CD4 T cell subset retains some proliferative ability, can respond to PD-1 blockade, and importantly gives rise to a “terminally exhausted” subset which is unresponsive to checkpoint blockade(52). Thus, these data interestingly suggest that in vivo anti-FAP CAR T therapy offers salutary effects via significant reduction of CD4 T cell exhaustion in MASH, and redirection of CD4 T cells away from the dysfunctional trajectory.

snRNAseq analysis revealed distinct hepatocyte clusters in each treatment group (Fig S8G, H). In comparing gene expression among hepatocyte clusters, there was broad downregulation of key fatty acid metabolism genes *Dgat2*, *Cpt1a*, *Scd1*, *Tm6sf2*, *Apoa4* and *Acot2* (Fig 6A-C), and downregulation of *Sox9* (Fig 6D), a key regulator of ECM deposition in the CD5-FAPCAR recipients, compared to the IgG-FAPCAR controls. Interestingly, *Cyp4a14* was the top DEG in CD5-FAPCAR hepatocytes compared IgG-FAPCAR hepatocytes (Fig 6E). Upregulation of cytochrome P450 omega-hydroxylase is linked to MASLD/MASH progression(34). This gene was expressed in **∼**80% of IgG-FAPCAR hepatocytes and MASH-saline hepatocytes but only expressed by ∼40% of CD5-FAPCAR hepatocytes, and ∼10% of Chow-saline hepatocytes (Fig 6F). Panther gene ontology analysis of the top, unique biological processes associated with the elevated DEGs in IgG-FAPCAR hepatocytes were heavily related to fatty acid metabolism, triglyceride synthesis, and insulin response (Fig 6G), consistent with a dysregulated hepatocyte phenotype seen in association with obesity, insulin resistance, dyslipidemia, and steatosis(53). On the other hand, biological processes associated with the elevated DEGs in CD5-FAPCAR hepatocytes were mostly related to nucleotide metabolism (Fig 6H), suggesting a recovery of physiological functions in hepatocytes.

**Fig. 6.**
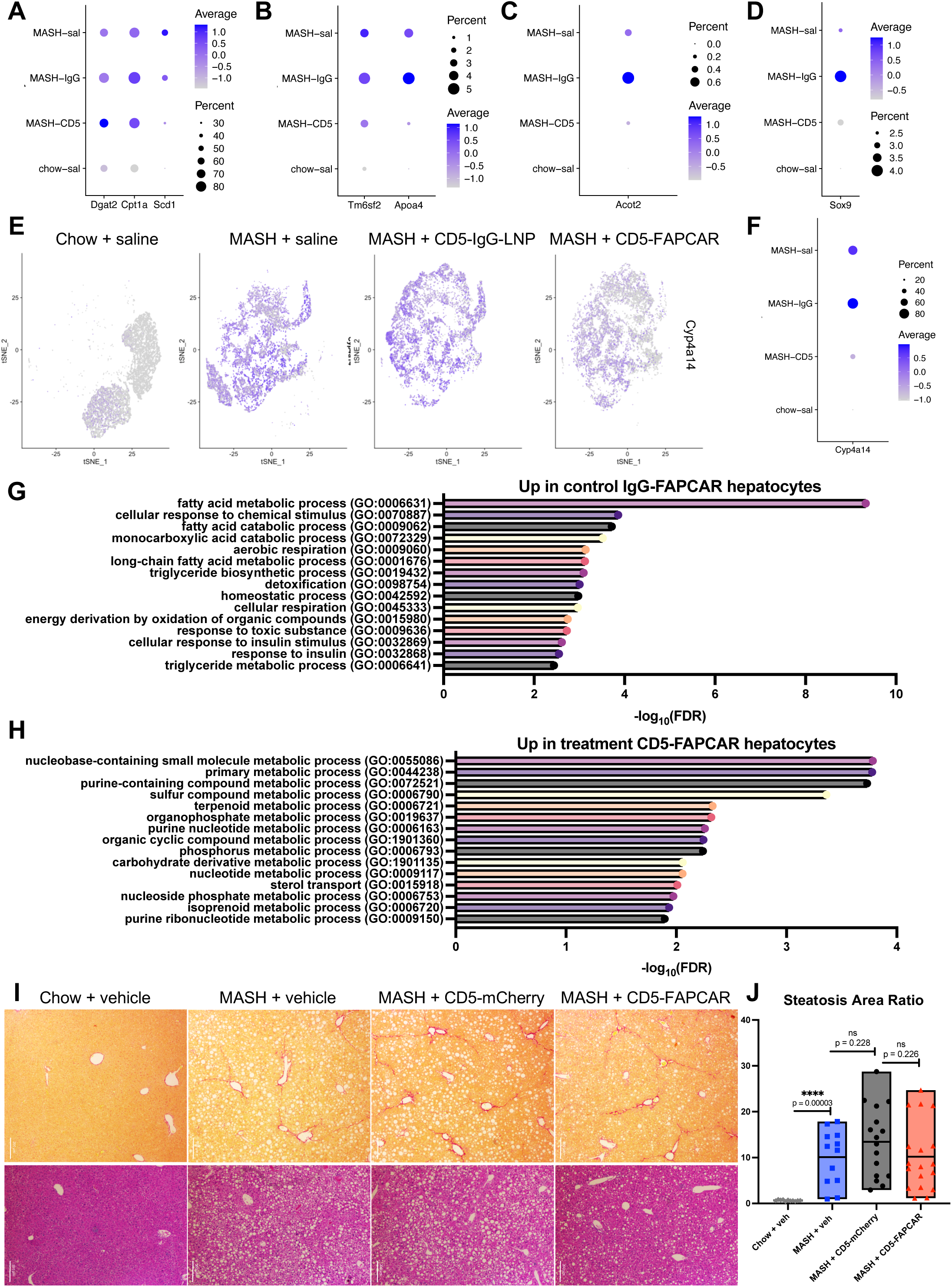
FAP^+^ HSC targeting via in vivo anti-FAP CAR T cells reprograms hepatocytes and promotes steatosis reduction. **(A-C)** DotPlots of RNA expression of key fatty acid metabolism genes and **(D)** *Sox9* in hepatocyte clusters, separated by experimental group. **(E)** FeaturePlots and **(F)** DotPlot of *Cyp4a14* expression in hepatocyte clusters, the top DEG between IgG- and CD5-FAPCAR groups. **(G)** Unique Panther Biological Processes associated with upregulated genes in IgG-FAPCAR control group. **(H)** Unique Panther Biological Processes associated with upregulated genes in CD5-FAPCAR treatment group. **(I)** Sirius Red (top) and H&E (bottom) representative images and **(J)** quantification steatosis area ratio via PharmaNest digital pathology analysis.

Steatosis analysis with digital microscopy (Fig 6I, J) reveals a trending reduction in hepatic steatosis in CD5-FAPCAR recipients, aligning with snRNAseq hepatocyte analysis and gene ontology data that suggest a resolution of aberrant lipid metabolism in CD5-FAPCAR-treated mice. These data suggest that depletion of FAP+ fibrogenic HSCs influences hepatocyte homeostasis and may contribute to MASH resolution through non-cell autonomous activities that restore healthy lipid metabolism, in part through attenuation of CYP4A14 activity.

### Anti-FAPCAR T therapy limits kidney fibrosis resulting from murine MASH

Advanced MASH and associated metabolic syndrome can lead to chronic kidney disease with renal fibrosis(54). The FAT MASH model recapitulates MASH-induced kidney injury(55), and FAP is expressed in injured kidney(56), thus we assessed the impact of in vivo anti-FAP CAR T therapy on renal fibrosis arising secondary to MASH. Sirius Red staining and digital microscopy revealed a trending reduction in fine and assembled collagen in the kidney cortex, and an overall trending reduction in the *Phenotypic Fibrosis Composite Score* of kidney cortex in CD5-FAPCAR mice (Fig S5A,B). These data suggest that in vivo-generated anti-FAP CAR T cells may also be efficacious towards renal disease associated with MASLD/MASH.

### In vivo anti-FAP CAR T cells are well-tolerated and transient

Thirty days after t-LNP injection there were no significant changes in AST, ALT, albumin, cholesterol or bilirubin (Fig S6A-F), or liver and spleen weights (Fig S6G, H). Throughout the 10 weeks of the FAT MASH model, including the 4 weeks post-t-LNP injection, there were no significant changes to body weight (Fig S6I). Flow cytometric analysis at the 30-day post-t-LNP timepoint revealed no FAPCAR+ immune cells remaining in the spleen or liver (Fig S6J). Together, these data highlight the transient, regulable activity of in vivo CAR T cells in clearing HSCs and reinforce the lack of hepatotoxicity.

## DISCUSSION

The growing appreciation of HSC heterogeneity allowed us to refine therapeutic targeting to only fibrogenic subsets while sparing quiescent HSCs. CAR T cells generated in vivo via t-LNP injection offer the prospect of regulated, self-limited depletion of pathogenic cells with little or no liver toxicity. We demonstrate that anti-CD5 t-LNPs delivering anti-FAPCAR mRNA are specifically taken up by splenic and hepatic CD5^+^ cells, generating FAPCAR^+^ T cells in vivo, which effectively deplete only fibrogenic HSC subsets, thereby directly reducing fibrosis in MASH. Furthermore, depletion of FAP^+^ HSCs promotes expansion of activated anti-fibrotic Ly6C^lo^ monocyte-derived macrophages, restores the healthy LSEC population, redirects CD4 T cells away from terminal exhaustion, and improves hepatocyte function, resulting in a reduction of lipid metabolism pathway related genes, an increase in pathways associated with normal hepatocyte function (purine synthesis pathways) and reduced *Cyp4a14* expression that is linked to hepatic steatosis and fibrosis.

*FAP* mRNA and protein is restricted to activated HSCs and their quiescent precursors in mouse and human MASH. Notably, significant FAP protein expression was observed in early MASH, aligning with earlier observations that FAP expression correlates with matrix metalloproteinases MMP1 and MMP13(57), at the advancing portion of cultured HSCs, to enhance their migration(58), collectively implicating FAP+ fibroblasts in fibrosis initiation and progression. These findings reinforce FAP as a potentially robust therapeutic target in MASH fibrosis(59-62).

By administering t-LNPs carrying FAPCAR mRNA coated with either anti-CD5 antibodies or IgG, we demonstrate their high specificity to CD5-expressing cells, generating FAPCAR T cells in spleen and liver within 18 hours of administration. Interestingly, we detected small numbers of other FAPCAR^+^ immune populations including macrophages, NKT cells, NK cells and B cells. While the intention is to generate CAR T cells, other kinds of CAR+ immune cells may be salutary as well. For example, CAR-NK, CAR-NKT and CAR-macrophage therapies are being developed that are potentially useful in cancer immunotherapies(63, 64). Notably, we observed generation of anti-FAP CAR macrophages in IgG-FAPCAR controls, likely through phagocytosis of IgG-coated t-LNPs (data not shown), which also had a subsequent trending reduction in fibrosis compared to MASH animals administered only saline. This finding aligns with recent data showing that anti-FAP CAR macrophages can moderately reduce chemically induced liver fibrosis(65). Thus, while the bulk of fibrosis resolution in anti-FAPCAR therapy is driven by CAR T cells, other CAR+ immune cells likely small contributions to fibrosis reduction as well.

Importantly, in vivo FAPCAR^+^ immune cells are cleared by four weeks, consistent earlier data demonstrating their presence for only 3-5 days(30). These reassuring findings reduce concern about potential unwanted effects of persistent, long-term in vivo CAR T cells. In contrast, ex vivo CAR T cells lead to DNA integration that allows for continued proliferation and expansion upon interaction with target antigen(66). Indeed, conventional CAR T used to treat hematologic malignancies may persist in patients for up to 10 years(33). Thus, in vivo CAR T-mediated therapy to deplete non-malignant cells such as HSCs may benefit from being self-limited, thereby avoiding disruption of liver homeostasis.

Fibrosis was significantly decreased four weeks after LNP injection, accompanied by reduced fibrogenic proteins and aSMA^+^ cells, and diminished *Fap* expression by snRNAseq. Flow cytometric tSNE analysis confirmed a relative reduction in activated HSCs and an expansion of quiescent HSCs, indicating that the in vivo anti-FAP CAR Ts are depleting FAP^+^ HSC subsets sufficiently to significantly reduce fibrosis. Intriguingly, snRNAseq and flow cytometry indicate a relative expansion of senescent HSCs in healthy chow-fed mice and in anti-FAPCAR t-LNP-treated MASH mice. The documented fluctuation of HSC senescence throughout the course of MASH(37) and their known profibrotic and antifibrotic(67) functions merit continued exploration.

Interestingly, non-HSC liver populations were affected in ways that would promote MASH resolution. The observed increases in activated “non-classical” Ly6c^lo^ monocyte-derived macrophages and LSECs would favor tissue repair and quell inflammation, thus tilting the scale towards fibrosis resolution and tissue recovery. Additionally, the observed reduction in PD-1^pos^ CD4 T cells, specifically in the progenitor exhausted CD4 T cell subset, likely contributes to limiting the progression of MASH, and may prevent future advancement to hepatocellular carcinoma(51). Although the exact mechanism of how FAP^+^ HSC depletion causes expansion of activated Ly6c^lo^ monocyte-derived macrophages and healthy LSECs and depletion of progenitor exhausted CD4 T cells, is currently unknown, these effects highlight the broad communication networks between HSCs and other cells in the hepatic milieu(36).

The striking reduction in fatty acid metabolism dysregulation in mice receiving in vivo anti-FAP CAR treatment, as indicated by differentially expressed genes including *Cyp4a14*, gene ontology pathway analysis, and a trending reduction in steatosis, suggests that FAP^+^ HSC depletion results in restored hepatocyte lipid homeostasis that may contribute to MASH resolution(34). Although the mechanism underlying improved lipid homeostasis is unknown, these striking changes hint at broader salutary effects of fibrogenic HSC subtype depletion on hepatocytes, which is further indicated by decreased *Sox9* expression.

The yin-yang relationship between fibrosis and hepatic regeneration is increasingly of interest, in which reduced fibrosis is linked to improved hepatocellular function. This relationship is evident in patients with advanced fibrosis treated effectively with antiviral therapies for HBV or HCV(68), or in a mouse study in which clearance of uPAR+ senescent HSCs in chronic liver disease led to increased serum albumin, a marker of hepatocyte health(10). Recent studies have uncovered paracrine signals that could support this relationship in liver(69) and kidney(70).

Intriguingly, our data also suggest that in vivo anti-FAP CAR T cells have salutary effects extending beyond the liver, leading to reduced renal cortical fibrosis which is common in MASH. Thus, targeting FAP via in vivo CAR T therapy may benefit both hepatic and extra-hepatic (i.e., renal) fibrosis from MASH.

Our proof-of-concept studies described here contain a few noteworthy limitations. First, the long-term efficacy and impact of in vivo CAR T therapy on liver regeneration and function are not yet established. Similarly, the durability of fibrosis reduction after one dose of in vivo CAR T, as well as the efficacy of in vivo CAR T cells in advanced MASH fibrosis are unknown. Additionally, the direct impact, if any, of LNPs on liver biology merits further evaluation, although the safety of LNP-based COVID-19 mRNA vaccines(71) is reassuring, with seemingly endless additional formulations under active development(72).

The value of in vivo CAR T therapy to clear fibrogenic cells in other liver diseases and tissues could be explored, as well. Further exploration of the cross talk between HSCs and other liver cells impacted by FAPCAR administration, and the mechanisms underlying effects on macrophage, endothelial, T cell and hepatocyte function and heterogeneity merit further clarification. Additional investigation of FAPCAR administration effect on kidney function in MASH may further reinforce the potential antifibrotic activity in multiple tissue pathologies associated with the disease(54).

In conclusion, these findings demonstrate that in murine MASH, in vivo anti-FAP CAR T cells directly improve hepatic fibrosis and indirectly restore hepatic homeostasis through non-cell autonomous effects on multiple cell types. Furthermore, our findings establish the importance of FAP^+^ HSCs in MASH fibrosis, and the potential benefit of in vivo anti-FAP CAR T cells that extends beyond clearance of fibrogenic cells.

## MATERIALS AND METHODS

### Animal models

Eight-week old male C57BL/6J mice purchased from Jackson Laboratory were placed on the Fibrosis and Tumor (FAT) MASH model, in which the mice are fed a Western diet (Teklad diets, 120528) of 21.1% fat, 41% sucrose, and 1.25% cholesterol and providing a sugar water solution containing 23.1g d-fructose/L and 18.9g d-glucose/L, combined with weekly intraperitoneal injections of CCl^4^ at the low dose of 0.2 ml (0.32 mg)/g of body weight(35). After six weeks of the FAT MASH model (or regular chow diet for controls), mice received tail vein injections (TVIs) of antibody-targeted lipid nanoparticles (t-LNPs) or vehicle administered in a total volume of 150uL. Two rounds of animal experiments were conducted. One involved TVI of 10ug of CD5 t-LNP carrying mCherry mRNA (CD5-mCherry), 10ug of CD5 t-LNP carrying anti-FAP CAR mRNA (CD5-FAPCAR) or vehicle (MilliQ water) per mouse. The other involved TVI of 10ug of IgG t-LNP carrying anti-FAP CAR mRNA (IgG-FAPCAR), 10ug of CD5 t-LNP carrying anti-FAP CAR mRNA (CD5-FAPCAR) or vehicle (saline) per mouse. A subset of mice was sacrificed 18 hours post-TVI, and the remaining mice were sacrificed 30 days post-TVI. Blood was drawn from the inferior vena cava with an aliquot stained for flow cytometry and the remainder used for serum analysis via VRL Diagnostics (Gaithersburg, Maryland, USA). Livers and spleens were weighed. A fragment of spleen was embedded in paraffin or OCT for histology, and the remainder was used for flow cytometry. The left liver lobe was sectioned and embedded in paraffin or OCT for histology, and 1g of liver tissue was used for flow cytometry. Remaining liver was minced and snap frozen for protein and RNA analyses.

### Targeted lipid nanoparticles

Targeted lipid nanoparticles (t-LNPs) containing mRNA encoding a chimeric antigen receptor (CAR) targeting fibroblast activation protein alpha (FAP) were generated by Capstan Therapeutics using an established protocol(30), and provided by Dr. Epstein at University of Pennsylvania. mRNA was generated by in vitro transcription with 1-methylpseudouridine. t-LNPs were shipped on dry ice and maintained at -80°C for 5 days. Thawed t-LNPs were immediately diluted with water or saline and injected via tail vein injection.

### Immunofluorescence for FAP, FAP + GFAP, aSMA

Formalin-fixed, paraffin-embedded liver sections were baked for one hour at 60°C, and deparaffinized with xylene followed by an ethanol gradient. Heat-induced epitope retrieval was done with sodium citrate buffer at pH 6.0 (Citrate Buffer, C9999) in a steamer for 30 minutes, followed by 30 minutes of cooling at room temperature. Slides were washed with 1x PBS, and blocked with Intercept blocking buffer (LICOR, #927-60001). Slides were incubated with primary antibody overnight at 4°C, all diluted in Intercept blocking buffer 1:200 (anti-FAP, Abcam ab218164; anti-GFAP, Abcam ab4674; anti-aSMA, Abcam ab5694). Slides were washed with 1x PBS, incubated with secondary antibodies at room temperature for one hour, all diluted in Intercept blocking buffer 1:500 (Alexa Fluor 647 donkey anti-rabbit, Invitrogen A31573; Alexa Fluor 555 goat anti-chicken IgG, Invitrogen A21437), washed with 1x PBS again, mounted with Fluoromount-G mounting media containing DAPI (Southern biotech, 0100-20), and imaged with Zeiss fluorescent microscope.

### Immunofluorescence for FAPCAR and lymphocyte markers

Formalin-fixed, paraffin-embedded liver sections were baked for one hour at 60°C, and deparaffinized with xylene followed by an ethanol gradient. Heat-induced epitope retrieval was done with sodium citrate buffer at pH 6.0 (Citrate Buffer, C9999) in a steamer for 30 minutes, followed by 30 minutes of cooling at room temperature. Slides were washed with 1x PBS, blocked with Intercept blocking buffer (LICOR, #927-60001), and washed with 1x PBS. Slides were blocked with avidin/biotin blocking kit (Vector Labs, SP-2001), and incubated overnight with biotin-conjugated goat anti-mouse IgG F(ab)^2^, (Jackson ImmunoResearch 115-065-072) mixed with anti-CD4 (Thermo Scientific 14-0041-82), anti-CD8a (Thermo Scientific, 14-0081-82) or anti-B220 (Thermo Scientific 14-0452-82), all 1:200 dilutions. Slides were washed with 1X PBS and incubated with Streptavidin Alexa Fluor 647 conjugate (Thermo Scientific, S32357) and Cy3-conjugated AffiniPure Donkey anti-rat IgG (Jackson ImmunoResearch 712-165-153) in Intercept blocking buffer, both at 1:500 dilutions. Slides were washed with 1x PBS and mounted with Fluoromount-G mounting media containing DAPI and imaged with Zeiss fluorescent microscope.

### Sirius Red Staining and AI-based collagen quantification in liver sections

Formalin-fixed, paraffin-embedded liver sections and kidney sections were baked for one hour at 60°C, and deparaffinized with xylene followed by an ethanol gradient, and a final wash in distilled water for two minutes. Slides were incubated in Sirius Red staining mix (Direct red 80 Sigma 365548-5G) for 1 hour in a shaker at room temperature, followed by three washes in distilled water, an ethanol dehydration gradient, and xylene. Slides were mounted with Permount mounting media (Fisher Scientific SP15500). Digital pathology analysis of fibrosis and steatosis was done using the FibroNest^TM^ platform. Sirius red-positive pixels were used to detect collagen. Tissue fibrosis phenotype was described for its collagen content and structure (‘assembled’ versus ‘fine’), the morphometric traits (shape and size) of the segmented collagen fibers, and fibrosis architecture traits. The fibrosis architecture analysis included collagen fiber disorganization, compactness, pattern presence, and relative density. Each fibrosis trait was quantified with seven statistical parameters to account for severity, progression, distortion, and variance for both the fine and assembled collagen fibers, resulting in up to 336 qFTs (quantitative Fibrosis Traits). This allowed for generation of composite scores for fibrosis severity for each phenotypic layer (collagen content, fibers morphometry, fibrosis architecture) and for the combined layers (Phenotypic-Fibrosis Composite Score (Ph-FCS)). Areas of macro-steatosis (vacuoles >7 mM diameter) were identified and distinguished from areas of vasculature and quantified to generate the Steatosis Area Ratio score.

### Single cell isolation and staining for flow cytometry

Blood: About 600uL of blood were collected per mouse via blood draw from the inferior vena cava immediately after cervical dislocation. About 150uL serum was collected and remaining blood was spun at 500g for 3 min, supernatant was aspirated, and cell pellet was resuspended in 150uL of 1x RBC lysis buffer (Thermo Fisher Scientific, A1049201) and incubated at room temperature for 3-5 minutes. 1350uL of 1X PBS was added, samples were pelleted, supernatant was aspirated, and sample was stained for flow cytometry.

Spleen: Spleen was bisected – half was used for histological analysis, and half was crushed through a 70u filter in a 50mL falcon tube, and washed with RPMI + 10% FBS until a total volume of 35mL was reached. Sample was spun at 370g for 5 minutes at 4°C, supernatant was dumped, and pellet was resuspended in 1mL of RBC lysis buffer for 3-5 minutes at room temperature. 30mL of PBS was added, sample was spun, and pellet was resuspended with 1mL and filtered with a 70u filter.

Liver: Hepatic non-parenchymal cells were isolated via an established protocol from murine MASH tissue(37). Antibodies used for flow cytometry staining are listed in Table S1. Flow gating strategies shown in Fig S7A-C. For flow tSNE analysis, 5 samples from each group were concatenated, selecting 5,000 CD45^-^ CD31^-^ Live Singlets from each sample, for a total of 25,000 cells per experimental group, and 100,000 total cells in the analysis. Heatmaps of aSMA, Ki-67, and CD206 expression were used to distinguish HSC subsets.

### Immunoblotting

Protein lysates were extracted from minced frozen liver with RIPA buffer (Thermo Fisher Scientific), and protein concentration was measured with the BCA assay. Proteins were separated by electrophoresis on 4–12% Bis-Tris gels (Invitrogen) and then transferred to 0.45μm PVDF membranes. The membranes were blocked for 1 hour at room temperature in Tris-buffered saline/0.1% Tween 20 (TBST) containing 5% non-fat milk and then incubated with primary antibody (anti-COL1A1, BIOSS BS-10423R 1:2,000; anti-αSMA, Abcam ab5694 1:2,000; anti-B-actin HRP conjugated, Cell Signaling 5125S 1:1,000) in Intercept blocking buffer at 4°C overnight. The membranes were then incubated with the corresponding HRP-conjugated secondary antibody (anti-rabbit IgG HRP 1:2,000 for COL1A1 and ACTA2) and proteins were detected with SuperSignal™ West Pico PLUS Chemiluminescent Substrate (Thermo Fisher Scientific).

### Nuclei isolation for snRNAseq sample submission

About 10-15mg of snap frozen liver tissue from four representative mice per group were pooled. Samples were prepared via an established protocol for nuclei isolation from murine MASH tissue(36). Solutions used included 2xST buffer (292 mM NaCl (Thermo Fisher Scientific, cat. no. AM9759)), 20 mM Tris-HCl pH 7.5 (Thermo Fisher Scientific, cat. no.15567027), 2 mM CaCl2 (VWR International Ltd, cat. no. 97062-820) and 42 mM MgCl2 (Sigma Aldrich, cat. no. 1028) in DNA/RNAse free water. 0.03% TST was prepared by adding 5mL of 2xST, 83uL Ribolock,83uL 2%BSA, 300uL of 1% Tween20, 4.6mL of H2O. The samples used for snRNAseq Chromium 3’ GEX were from snap frozen Chow + saline, MASH + saline, MASH + IgG-FAPCAR, and MASH + CD5-FAPCAR livers, about 3 months after euthanasia and cryopreservation. Each sample was chopped for 10 minutes over ice and passed through a strainer with 1mL TST. Samples were spun down at 500xg for 5 minutes at 4oC, supernatant was aspirated. Pellet was resuspended by pipetting 30 times with P1000 tips in 200uL of 0.4% BSA/PBS, and then brought to the Genetics Core.

### snRNAseq analysis

snRNAseq data quality control was performed with CellBender to remove ambient RNA. Seurat pipeline was used for downstream clustering and analysis. Dimensional reduction was performed with the top 2000 highly variable genes. A mitochondrial cutoff of 5% and nFeature range of 200 to 5500 were applied. The top 20 principal components were used to perform principal component analysis, and clustering was performed using a resolution of 0.5. HSC cluster was scaled and subclustered using principal component (PC) analysis reduction with the top 30 principal components, top 2000 highly variable features, and with a resolution of 0.5. Hepatocyte cluster and endothelial cell cluster were scaled and subclustered using principal component (PC) analysis reduction with the top 30 principal components, top 2000 highly variable features, and with a resolution of 0.1. Endothelial cell cluster was further stratified by setting a cutoff of >3.5 *Ptprb* expression to remove contaminating hepatocytes. Differentially expressed gene (DEG) analysis was performed between the IgG-FAPCAR and CD5-FAPCAR groups. DEGs with adjusted p values < 1x10^-5^ were considered significant. Significant hepatocyte DEGs were input into the AmiGO2 web server (http://amigo.geneontology.org/amigo). The top 15 Panther biological processes that were significantly enriched pathways (FDR <0.05) with more than one gene reported were included.

### Statistical analysis

Results are shown as means ± standard deviation. Statistical analysis was performed using Student’s t test in Prism 9.

### List of Supplementary Materials

Fig S1 to S8

Table S1

## Acknowledgements

This work was supported in part through the flow cytometry CoRE and the biorepository & pathology CoRE resources of the Tisch Cancer Institute.

This work was supported in part through the computational and data resources and staff expertise provided by Scientific Computing and Data at the Icahn School of Medicine at Mount Sinai and by the Clinical and Translational Science Awards (CTSA) grant UL1TR004419 from the National Center for Advancing Translational Sciences. Research reported in this publication was also supported by the Office of Research Infrastructure of the National Institutes of Health under award number S10OD026880 and S10OD030463. The content is solely the responsibility of the authors and does not represent the official views of the National Institutes of Health.

## Funding

CNY: T32GM007280, the Medical Scientist Training Program Training Grant and NCI T32CA078207-22, the NCI-funded Training Program in Cancer Biology

KL: T32GM007280, the Medical Scientist Training Program Training Grant and NCI T32 CA 078207-24 the NCI-funded Training Program in Cancer Biology

SW: 1R01 DK136016-01

JGR: Swedish Cancer Foundation, Cancer Research Karolinska Institutet, and the Sjöberg Foundation.

JAE: Sponsored research agreement from Capstan Therapeutics, R35 HL140018, the Cotswold Foundation.

SLF: 5R01DK128289-03, 5R01 DK121154-04, 5P30CA196521-08; UL1TR004419

## Author contributions

Conceptualization: HA, JGR, JAE, SLF

Data curation: CNY

Formal analysis: CNY

Funding acquisition: CNY, JAE, SLF

Investigation: CNY, BC, TQ, TT

Methodology: CNY, TW, TEP, RR, LC, AL, HA, HP, JAE

Project administration: CNY, BC, SLF

Resources: TW, TEP, RR, LC, AL, HA, HP, SW, JAE, SLF

Software: CNY, KL

Supervision: SW, JGR, JAE, SLF

Validation: CNY, BC, TQ, TT

Visualization: CNY, BC, TT, RR, LC, AL

Writing – original draft: CNY, SLF

Writing – review & editing: CNY, BC, LC, SW, JGR, JAE, SLF

## Data and materials availability

snRNAseq data will be made available on Gene Expression Omnibus repository

## References and Notes

1. Miao L, Targher G, Byrne CD, Cao YY, Zheng MH. Current status and future trends of the global burden of MASLD. Trends Endocrinol Metab. 2024.

2. Younossi ZM, Golabi P, Paik JM, Henry A, Van Dongen C, Henry L. The global epidemiology of nonalcoholic fatty liver disease (NAFLD) and nonalcoholic steatohepatitis (NASH): a systematic review. Hepatology. 2023;77(4):1335–47.

3. Younossi ZM. The epidemiology of nonalcoholic steatohepatitis. Clin Liver Dis (Hoboken). 2018;11(4):92–4.

4. Anstee QM, Targher G, Day CP. Progression of NAFLD to diabetes mellitus, cardiovascular disease or cirrhosis. Nat Rev Gastroenterol Hepatol. 2013;10(6):330–44.

5. Friedman SL, Neuschwander-Tetri BA, Rinella M, Sanyal AJ. Mechanisms of NAFLD development and therapeutic strategies. Nat Med. 2018;24(7):908–22.

6. Wang X, Zhang L, Dong B. Molecular mechanisms in MASLD/MASH related HCC. Hepatology. 2024.

7. Wang S, Friedman SL. Found in translation-Fibrosis in metabolic dysfunction-associated steatohepatitis (MASH). Sci Transl Med. 2023;15(716):eadi0759.

8. Schuppan D, Kim YO. Evolving therapies for liver fibrosis. J Clin Invest. 2013;123(5):1887–901.

9. Taylor RS, Taylor RJ, Bayliss S, Hagstrom H, Nasr P, Schattenberg JM, et al. Association Between Fibrosis Stage and Outcomes of Patients With Nonalcoholic Fatty Liver Disease: A Systematic Review and Meta-Analysis. Gastroenterology. 2020;158(6):1611–25 e12.

10. Amor C, Feucht J, Leibold J, Ho YJ, Zhu C, Alonso-Curbelo D, et al. Senolytic CAR T cells reverse senescence-associated pathologies. Nature. 2020;583(7814):127-32.

11. Geerts A. History, heterogeneity, developmental biology, and functions of quiescent hepatic stellate cells. Semin Liver Dis. 2001;21(3):311–35.

12. Eng FJ, Friedman SL. Fibrogenesis I. New insights into hepatic stellate cell activation: the simple becomes complex. Am J Physiol Gastrointest Liver Physiol. 2000;279(1):G7–G11.

13. Tsuchida T, Friedman SL. Mechanisms of hepatic stellate cell activation. Nat Rev Gastroenterol Hepatol. 2017;14(7):397–411.

14. Koyama Y, Brenner DA. Liver inflammation and fibrosis. J Clin Invest. 2017;127(1):55–64.

15. Cogliati B, Yashaswini CN, Wang S, Sia D, Friedman SL. Friend or foe? The elusive role of hepatic stellate cells in liver cancer. Nat Rev Gastroenterol Hepatol. 2023;20(10):647–61.

16. Kisseleva T, Brenner D. Molecular and cellular mechanisms of liver fibrosis and its regression. Nat Rev Gastroenterol Hepatol. 2021;18(3):151–66.

17. Rosenthal SB, Liu X, Ganguly S, Dhar D, Pasillas MP, Ricciardelli E, et al. Heterogeneity of HSCs in a Mouse Model of NASH. Hepatology. 2021;74(2):667–85.

18. Kostallari E, Wei B, Sicard D, Li J, Cooper SA, Gao J, et al. Stiffness is associated with hepatic stellate cell heterogeneity during liver fibrosis. Am J Physiol Gastrointest Liver Physiol. 2022;322(2):G234–G46.

19. Cheng S, Zou Y, Zhang M, Bai S, Tao K, Wu J, et al. Single-cell RNA sequencing reveals the heterogeneity and intercellular communication of hepatic stellate cells and macrophages during liver fibrosis. MedComm (2020). 2023;4(5):e378.

20. Trinh VQ, Lee TF, Lemoinne S, Ray KC, Ybanez MD, Tsuchida T, et al. Hepatic stellate cells maintain liver homeostasis through paracrine neurotrophin-3 signaling that induces hepatocyte proliferation. Sci Signal. 2023;16(787):eadf6696.

21. Kawahara A, Kanno K, Yonezawa S, Otani Y, Kobayashi T, Tazuma S, et al. Depletion of hepatic stellate cells inhibits hepatic steatosis in mice. J Gastroenterol Hepatol. 2022;37(10):1946–54.

22. Levy MT, McCaughan GW, Abbott CA, Park JE, Cunningham AM, Muller E, et al. Fibroblast activation protein: a cell surface dipeptidyl peptidase and gelatinase expressed by stellate cells at the tissue remodelling interface in human cirrhosis. Hepatology. 1999;29(6):1768–78.

23. Levy MT, McCaughan GW, Marinos G, Gorrell MD. Intrahepatic expression of the hepatic stellate cell marker fibroblast activation protein correlates with the degree of fibrosis in hepatitis C virus infection. Liver. 2002;22(2):93–101.

24. Fan MH, Zhu Q, Li HH, Ra HJ, Majumdar S, Gulick DL, et al. Fibroblast Activation Protein (FAP) Accelerates Collagen Degradation and Clearance from Lungs in Mice. J Biol Chem. 2016;291(15):8070–89.

25. Yang AT, Kim YO, Yan XZ, Abe H, Aslam M, Park KS, et al. Fibroblast Activation Protein Activates Macrophages and Promotes Parenchymal Liver Inflammation and Fibrosis. Cell Mol Gastroenterol Hepatol. 2023;15(4):841–67.

26. Falamarzi K, Malekpour M, Tafti MF, Azarpira N, Behboodi M, Zarei M. The role of FGF21 and its analogs on liver associated diseases. Front Med (Lausanne). 2022;9:967375.

27. Harrison SA, Rolph T, Knott M, Dubourg J. FGF21 agonists: An emerging therapeutic for metabolic dysfunction-associated steatohepatitis and beyond. J Hepatol. 2024;81(3):562–76.

28. Schuster SJ, Bishop MR, Tam CS, Waller EK, Borchmann P, McGuirk JP, et al. Tisagenlecleucel in Adult Relapsed or Refractory Diffuse Large B-Cell Lymphoma. N Engl J Med. 2019;380(1):45–56.

29. June CH, Sadelain M. Chimeric Antigen Receptor Therapy. N Engl J Med. 2018;379(1):64–73.

30. Rurik JG, Tombacz I, Yadegari A, Mendez Fernandez PO, Shewale SV, Li L, et al. CAR T cells produced in vivo to treat cardiac injury. Science. 2022;375(6576):91-6.

31. Aghajanian H, Kimura T, Rurik JG, Hancock AS, Leibowitz MS, Li L, et al. Targeting cardiac fibrosis with engineered T cells. Nature. 2019;573(7774):430-3.

32. Ellebrecht CT, Bhoj VG, Nace A, Choi EJ, Mao X, Cho MJ, et al. Reengineering chimeric antigen receptor T cells for targeted therapy of autoimmune disease. Science. 2016;353(6295):179-84.

33. Melenhorst JJ, Chen GM, Wang M, Porter DL, Chen C, Collins MA, et al. Decade-long leukaemia remissions with persistence of CD4(+) CAR T cells. Nature. 2022;602(7897):503-9.

34. Zhang X, Li S, Zhou Y, Su W, Ruan X, Wang B, et al. Ablation of cytochrome P450 omega-hydroxylase 4A14 gene attenuates hepatic steatosis and fibrosis. Proc Natl Acad Sci U S A. 2017;114(12):3181–5.

35. Tsuchida T, Lee YA, Fujiwara N, Ybanez M, Allen B, Martins S, et al. A simple diet- and chemical-induced murine NASH model with rapid progression of steatohepatitis, fibrosis and liver cancer. J Hepatol. 2018;69(2):385–95.

36. Wang S, Li K, Pickholz E, Dobie R, Matchett KP, Henderson NC, et al. An autocrine signaling circuit in hepatic stellate cells underlies advanced fibrosis in nonalcoholic steatohepatitis. Sci Transl Med. 2023;15(677):eadd3949.

37. Yashaswini CN, Qin T, Bhattacharya D, Amor C, Lowe S, Lujambio A, et al. Phenotypes and ontogeny of senescent hepatic stellate cells in metabolic dysfunction-associated steatotic liver disease. J Hepatol. 2024.

38. Pfister D, Nunez NG, Pinyol R, Govaere O, Pinter M, Szydlowska M, et al. NASH limits anti-tumour surveillance in immunotherapy-treated HCC. Nature. 2021;592(7854):450-6.

39. Kostadinova R, Strobel S, Chen L, Fiaschetti-Egli K, Gadient J, Pawlowska A, et al. Digital pathology with artificial intelligence analysis provides insight to the efficacy of anti-fibrotic compounds in human 3D MASH model. Sci Rep. 2024;14(1):5885.

40. March S, Graupera M, Rosa Sarrias M, Lozano F, Pizcueta P, Bosch J, et al. Identification and functional characterization of the hepatic stellate cell CD38 cell surface molecule. Am J Pathol. 2007;170(1):176–87.

41. Alkhani A, Levy CS, Tsui M, Rosenberg KA, Polovina K, Mattis AN, et al. Ly6c(Lo) non-classical monocytes promote resolution of rhesus rotavirus-mediated perinatal hepatic inflammation. Sci Rep. 2020;10(1):7165.

42. Ramachandran P, Pellicoro A, Vernon MA, Boulter L, Aucott RL, Ali A, et al. Differential Ly-6C expression identifies the recruited macrophage phenotype, which orchestrates the regression of murine liver fibrosis. Proc Natl Acad Sci U S A. 2012;109(46):E3186–95.

43. Thomas G, Tacke R, Hedrick CC, Hanna RN. Nonclassical patrolling monocyte function in the vasculature. Arterioscler Thromb Vasc Biol. 2015;35(6):1306–16.

44. Nasiri-Ansari N, Androutsakos T, Flessa CM, Kyrou I, Siasos G, Randeva HS, et al. Endothelial Cell Dysfunction and Nonalcoholic Fatty Liver Disease (NAFLD): A Concise Review. Cells. 2022;11(16).

45. Wilson C, Lee MD, Buckley C, Zhang X, McCarron JG. Mitochondrial ATP Production is Required for Endothelial Cell Control of Vascular Tone. Function (Oxf). 2023;4(2):zqac063.

46. Eelen G, Dubois C, Cantelmo AR, Goveia J, Bruning U, DeRan M, et al. Role of glutamine synthetase in angiogenesis beyond glutamine synthesis. Nature. 2018;561(7721):63-9.

47. Xie Y, Lin H, Wei W, Kong Y, Fang Q, Chen E, et al. LINC00839 promotes malignancy of liver cancer via binding FMNL2 under hypoxia. Sci Rep. 2022;12(1):18757.

48. Su T, Qin XY, Dohmae N, Wei F, Furutani Y, Kojima S, et al. Inhibition of Ganglioside Synthesis Suppressed Liver Cancer Cell Proliferation through Targeting Kinetochore Metaphase Signaling. Metabolites. 2021;11(3).

49. Jung SY, Lee HK, Kim H, Kim S, Kim JS, Kang JG, et al. Depletion of ST6GALNACIII retards A549 non-small cell lung cancer cell proliferation by downregulating transferrin receptor protein 1 expression. Biochem Biophys Res Commun. 2021;575:78–84.

50. Nowak-Sliwinska P, van Beijnum JR, Huijbers EJM, Gasull PC, Mans L, Bex A, et al. Oncofoetal insulin receptor isoform A marks the tumour endothelium; an underestimated pathway during tumour angiogenesis and angiostatic treatment. Br J Cancer. 2019;120(2):218–28.

51. Sim BC, Kang YE, You SK, Lee SE, Nga HT, Lee HY, et al. Hepatic T-cell senescence and exhaustion are implicated in the progression of fatty liver disease in patients with type 2 diabetes and mouse model with nonalcoholic steatohepatitis. Cell Death Dis. 2023;14(9):618.

52. Miggelbrink AM, Jackson JD, Lorrey SJ, Srinivasan ES, Waibl-Polania J, Wilkinson DS, et al. CD4 T-Cell Exhaustion: Does It Exist and What Are Its Roles in Cancer? Clin Cancer Res. 2021;27(21):5742–52.

53. Sookoian S, Pirola CJ, Valenti L, Davidson NO. Genetic Pathways in Nonalcoholic Fatty Liver Disease: Insights From Systems Biology. Hepatology. 2020;72(1):330–46.

54. Sandireddy R, Sakthivel S, Gupta P, Behari J, Tripathi M, Singh BK. Systemic impacts of metabolic dysfunction-associated steatotic liver disease (MASLD) and metabolic dysfunction-associated steatohepatitis (MASH) on heart, muscle, and kidney related diseases. Front Cell Dev Biol. 2024;12:1433857.

55. Li X, Bhattacharya D, Yuan Y, Wei C, Zhong F, Ding F, et al. Chronic kidney disease in a murine model of non-alcoholic steatohepatitis (NASH). Kidney Int. 2024;105(3):540–61.

56. Byun JW, Paeng JC, Kim YJ, Lee SP, Lee YS, Choi H, et al. Evaluation of Fibroblast Activation Protein Expression Using (68)Ga-FAPI46 PET in Hypertension-Induced Tissue Changes. J Nucl Med. 2024;65(11):1776–81.

57. Bauer S, Jendro MC, Wadle A, Kleber S, Stenner F, Dinser R, et al. Fibroblast activation protein is expressed by rheumatoid myofibroblast-like synoviocytes. Arthritis Res Ther. 2006;8(6):R171.

58. Wang XM, Yu DM, McCaughan GW, Gorrell MD. Fibroblast activation protein increases apoptosis, cell adhesion, and migration by the LX-2 human stellate cell line. Hepatology. 2005;42(4):935–45.

59. Basalova N, Alexandrushkina N, Grigorieva O, Kulebyakina M, Efimenko A. Fibroblast Activation Protein Alpha (FAPalpha) in Fibrosis: Beyond a Perspective Marker for Activated Stromal Cells? Biomolecules. 2023;13(12).

60. Yang P, Luo Q, Wang X, Fang Q, Fu Z, Li J, et al. Comprehensive Analysis of Fibroblast Activation Protein Expression in Interstitial Lung Diseases. Am J Respir Crit Care Med. 2023;207(2):160–72.

61. Adams TS, Schupp JC, Poli S, Ayaub EA, Neumark N, Ahangari F, et al. Single-cell RNA-seq reveals ectopic and aberrant lung-resident cell populations in idiopathic pulmonary fibrosis. Sci Adv. 2020;6(28):eaba1983.

62. Lake BB, Menon R, Winfree S, Hu Q, Melo Ferreira R, Kalhor K, et al. An atlas of healthy and injured cell states and niches in the human kidney. Nature. 2023;619(7970):585-94.

63. Pan K, Farrukh H, Chittepu V, Xu H, Pan CX, Zhu Z. CAR race to cancer immunotherapy: from CAR T, CAR NK to CAR macrophage therapy. J Exp Clin Cancer Res. 2022;41(1):119.

64. Hadiloo K, Tahmasebi S, Esmaeilzadeh A. CAR-NKT cell therapy: a new promising paradigm of cancer immunotherapy. Cancer Cell Int. 2023;23(1):86.

65. Mao Y, Yao C, Zhang S, Zeng Q, Wang J, Sheng C, et al. Targeting fibroblast activation protein with chimeric antigen receptor macrophages. Biochem Pharmacol. 2024;230(Pt 3):116604.

66. Sterner RC, Sterner RM. CAR-T cell therapy: current limitations and potential strategies. Blood Cancer J. 2021;11(4):69.

67. Zhang M, Serna-Salas S, Damba T, Borghesan M, Demaria M, Moshage H. Hepatic stellate cell senescence in liver fibrosis: Characteristics, mechanisms and perspectives. Mech Ageing Dev. 2021;199:111572.

68. Rockey DC, Friedman SL. Fibrosis Regression After Eradication of Hepatitis C Virus: From Bench to Bedside. Gastroenterology. 2021;160(5):1502–20 e1.

69. Wang P, Kang Q, Wu WS, Rui L. Hepatic Snai1 and Snai2 promote liver regeneration and suppress liver fibrosis in mice. Cell Rep. 2024;43(3):113875.

70. Aggarwal S, Wang Z, Rincon Fernandez Pacheco D, Rinaldi A, Rajewski A, Callemeyn J, et al. SOX9 switch links regeneration to fibrosis at the single-cell level in mammalian kidneys. Science. 2024;383(6685):eadd6371.

71. Klein NP, Lewis N, Goddard K, Fireman B, Zerbo O, Hanson KE, et al. Surveillance for Adverse Events After COVID-19 mRNA Vaccination. JAMA. 2021;326(14):1390–9.

72. Hou X, Zaks T, Langer R, Dong Y. Lipid nanoparticles for mRNA delivery. Nat Rev Mater. 2021;6(12):1078–94.

