## Supplemental Figure 1 for "In vivo anti-FAP CAR T therapy reduces fibrosis and restores liver homeostasis in metabolic dysfunction-associated steatohepatitis"

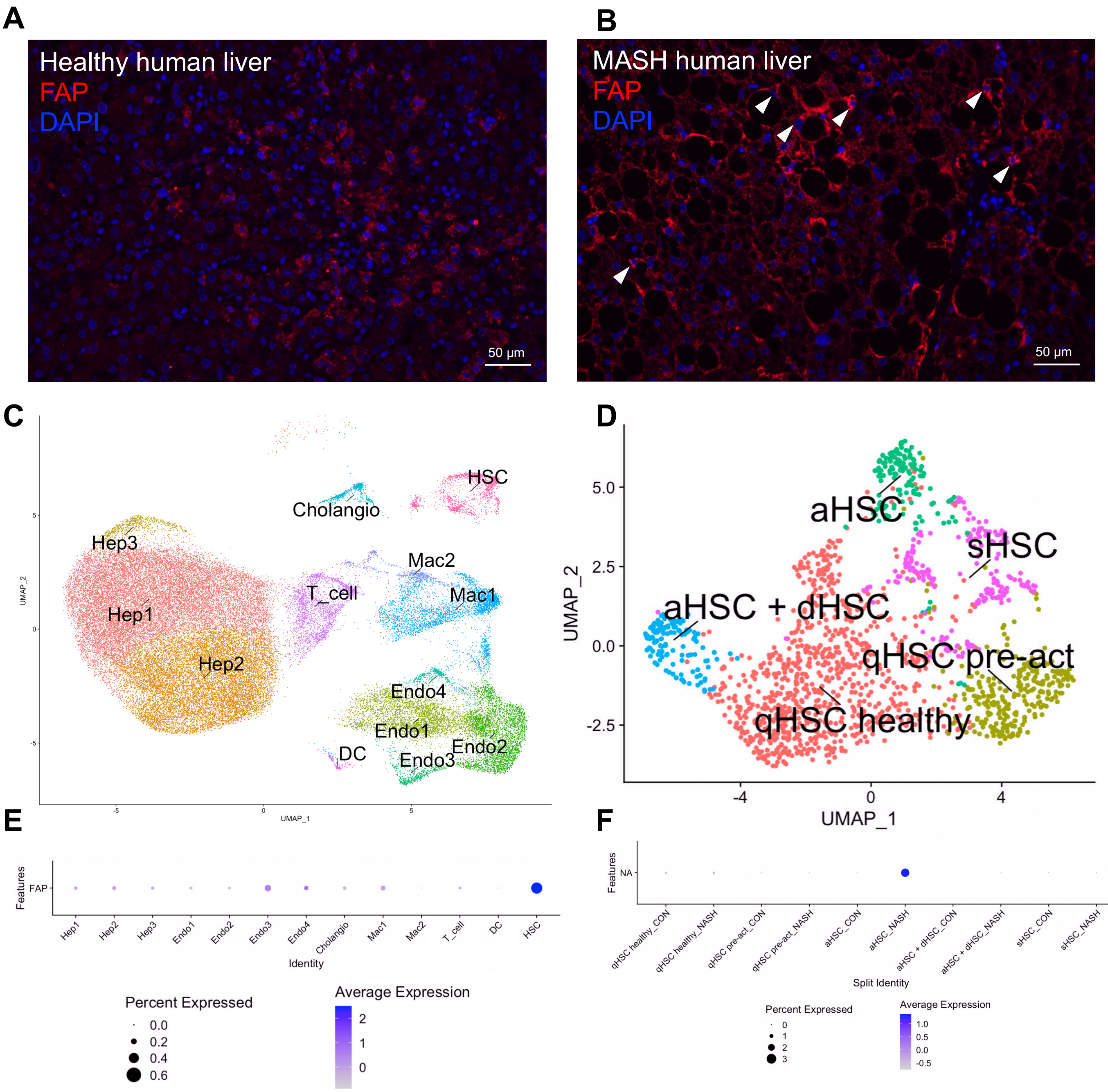


**Fig. S1. FAP is expressed by activated HSCs in human MASH. (A)** FAP immunostaining in normal human liver tissue and **(B)** MASH human liver tissue with FAP in red, DAPI in blue, and white arrowheads indicated FAP$^{+}$ cells. **(C)** snRNAseq Dimensional Plot of human MASH liver. **(D)** snRNAseq Dimensional Plot of HSC subsets in human MASH liver. **(E)** Dot Plot of FAP expression in human MASH liver. **(F)** Dot Plot of FAP expression in HSC subsets of human MASH liver.
