## Supplemental Figure 2 for "In vivo anti-FAP CAR T therapy reduces fibrosis and restores liver homeostasis in metabolic dysfunction-associated steatohepatitis"

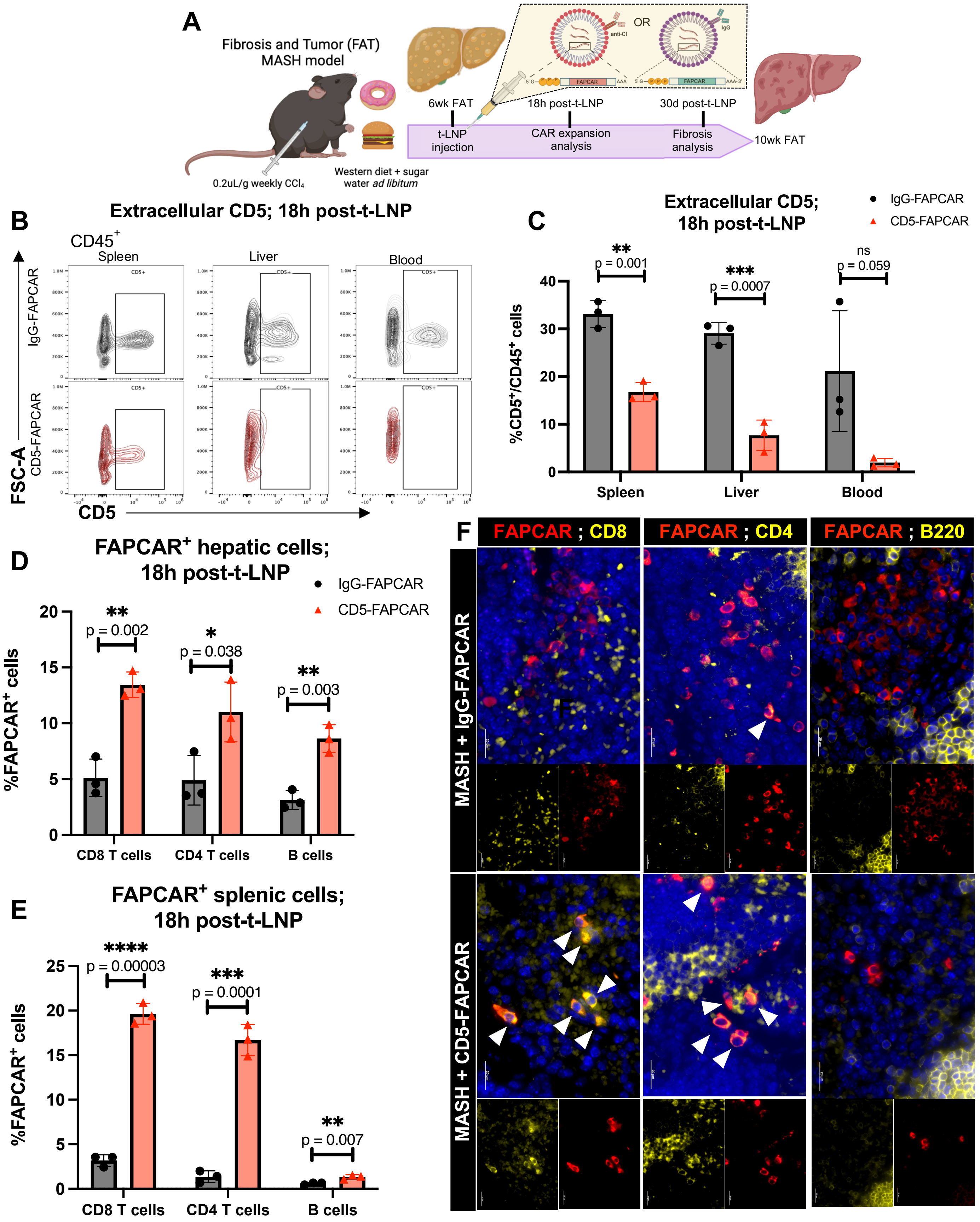

A
Fibrosis and Tumor (FAT) MASH model
0.2uL/g weekly CCl4
Western diet + sugar water ad libitum
6wk FAT
18h post-t-LNP
30d post-t-LNP
t-LNP injection
CAR expansion analysis
Fibrosis analysis
10wk FAT
OR
B
Extracellular CD5; 18h post-t-LNP
CD45+
Spleen
Liver
Blood
IgG-FAPCAR
CD5-FAPCAR
FSC-A
CD5
C
Extracellular CD5; 18h post-t-LNP
IgG-FAPCAR
CD5-FAPCAR
%CD5+/CD45+ cells
p = 0.001
p = 0.0007
ns p = 0.059
Spleen
Liver
Blood
D
FAPCAR+ hepatic cells; 18h post-t-LNP
%FAPCAR+ cells
p = 0.002
p = 0.038
p = 0.003
CD8 T cells
CD4 T cells
B cells
E
FAPCAR+ splenic cells; 18h post-t-LNP
%FAPCAR+ cells
p = 0.00003
p = 0.0001
p = 0.007
CD8 T cells
CD4 T cells
B cells
F
FAPCAR ; CD8
FAPCAR ; CD4
FAPCAR ; B220
MASH + IgG-FAPCAR
MASH + CD5-FAPCAR

**Fig. S2. CD5-targeted LNPs generate FAPCAR T cells in vivo. (A)** Experimental setup where treatment is CD5-FAPCAR t-LNP and control is IgG-FAPCAR t-LNP. **(B)** Flow plots and **(C)** quantification of extracellular CD5 levels in $CD45^+$ cells in IgG-FAPCAR control (black) vs CD5-FAPCAR treatment (red) in spleen, liver and blood, 18 hours after t-LNP injection. **(D)** Flow quantification of hepatic and **(E)** splenic FAPCAR+ lymphocytes. **(F)** Co-immunostaining of FAPCAR (red), respective lymphocyte markers (yellow) and DAPI (blue) in IgG-FAPCAR control spleen (top) and CD5-FAPCAR treatment spleen (bottom), where white arrows indicate areas of co-staining.
