## Supplemental Figure 3 for "In vivo anti-FAP CAR T therapy reduces fibrosis and restores liver homeostasis in metabolic dysfunction-associated steatohepatitis"

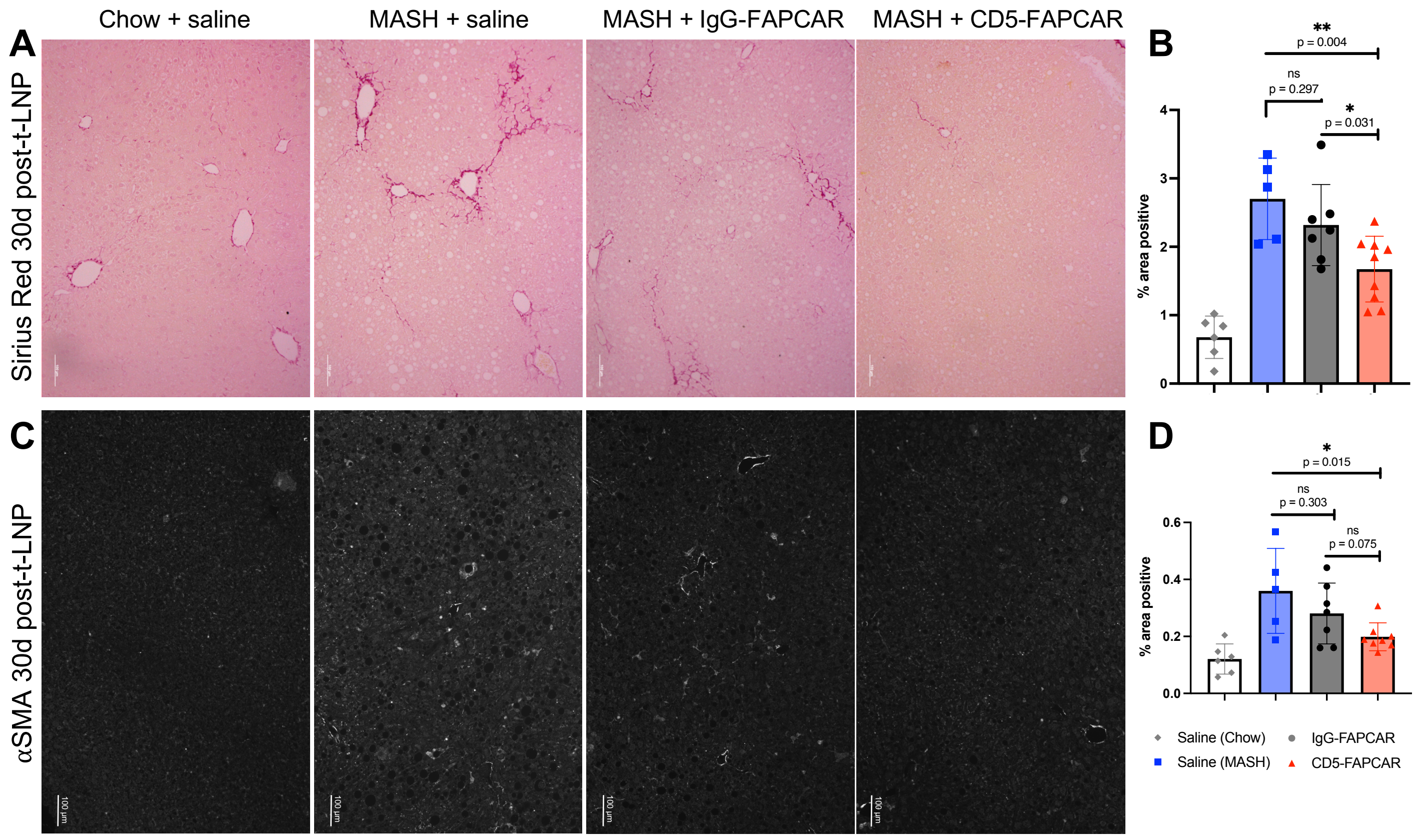


**Fig. S3. in vivo anti-FAP CAR T cells deplete activated HSCs and reduce hepatic fibrosis in MASH. (A)** Sirius Red staining with **(B)** quantification and **(C)** αSMA staining with **(D)** quantification of IgG-FAPCAR vs CD5-FAPCAR 30 days after t-LNP injection.
