## Supplemental Figure 4 for "In vivo anti-FAP CAR T therapy reduces fibrosis and restores liver homeostasis in metabolic dysfunction-associated steatohepatitis"

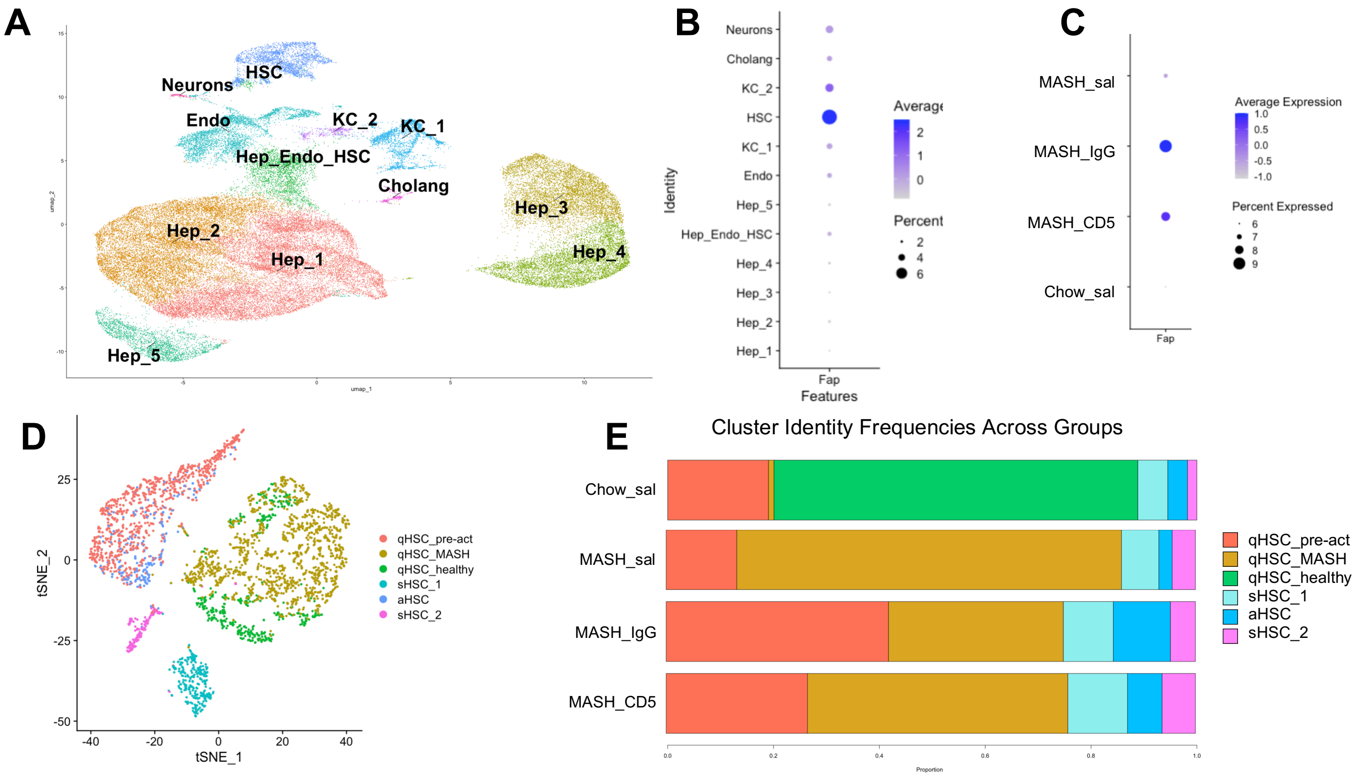


**Fig. S4. in vivo anti-FAP CAR T cells reduce FAP expression and cause redistribution of HSC subsets. (A)** snRNAseq Dimensional Plot and **(B)** Dot Plot of FAP expression of different hepatic cell populations. **(C)** Dot Plot of FAP expression in each experimental group. **(D)** snRNAseq Dimensional Plot of HSC subsets. **(E)** Bar Plots of relative proportions of HSC subsets. All snRNAseq data are from Chow + saline, MASH + saline, MASH + IgG-FAPCAR and MASH + CD5-FAPCAR experiment.
