## Supplemental Figure 5 for "In vivo anti-FAP CAR T therapy reduces fibrosis and restores liver homeostasis in metabolic dysfunction-associated steatohepatitis"

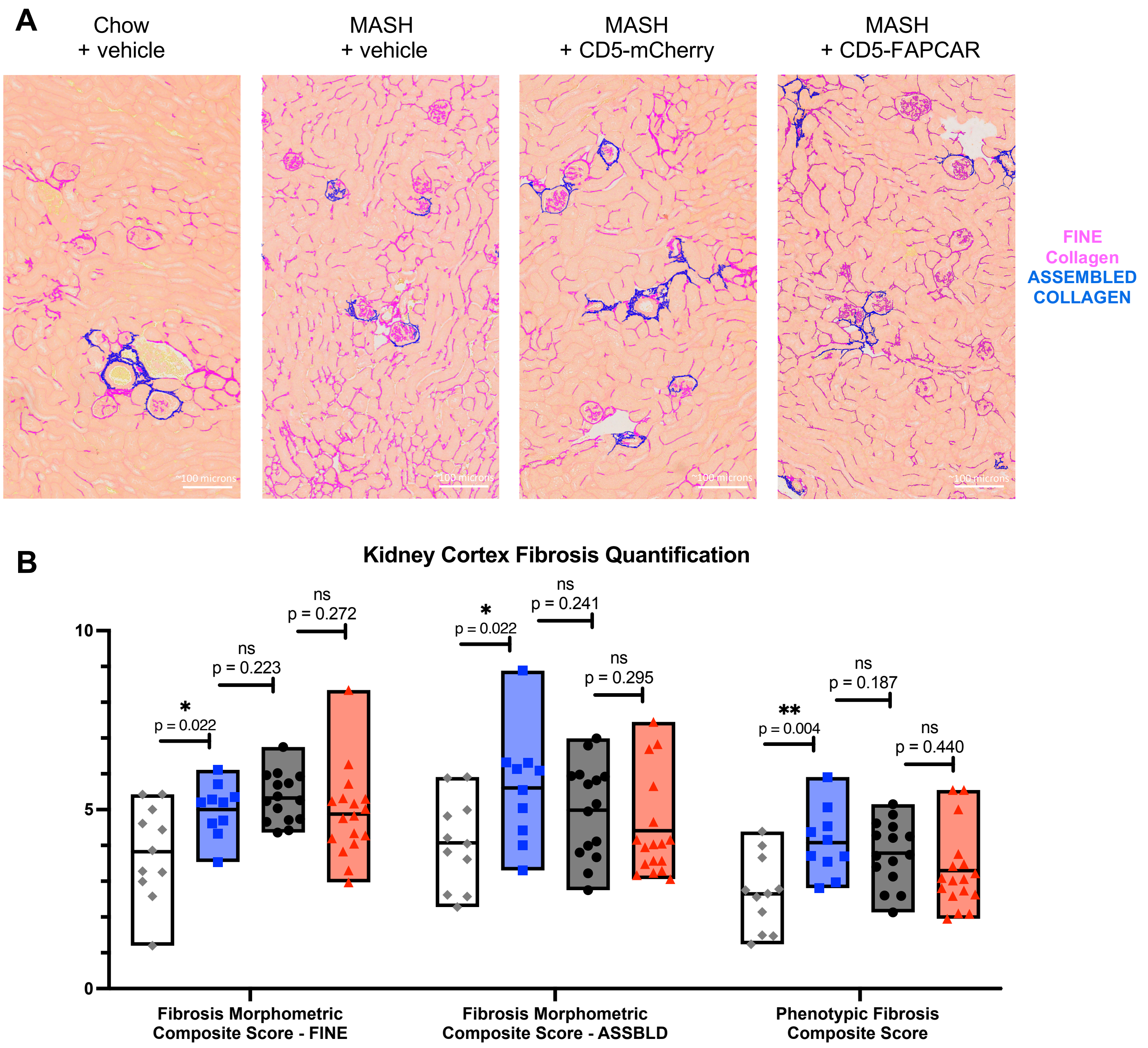


**Fig. S5. in vivo anti-FAP CAR T cells reduce renal fibrosis in MASH. (A)** Digital pathology analysis of renal cortex via FibroNest$^{TM}$ platform of fine collagen (pink) and assembled collagen (blue), overlaid on Sirius Red. **(B)** Digital pathology quantification of renal cortex fibrosis severity via FibroNest$^{TM}$.
