## Supplemental Figure 6 for "In vivo anti-FAP CAR T therapy reduces fibrosis and restores liver homeostasis in metabolic dysfunction-associated steatohepatitis"

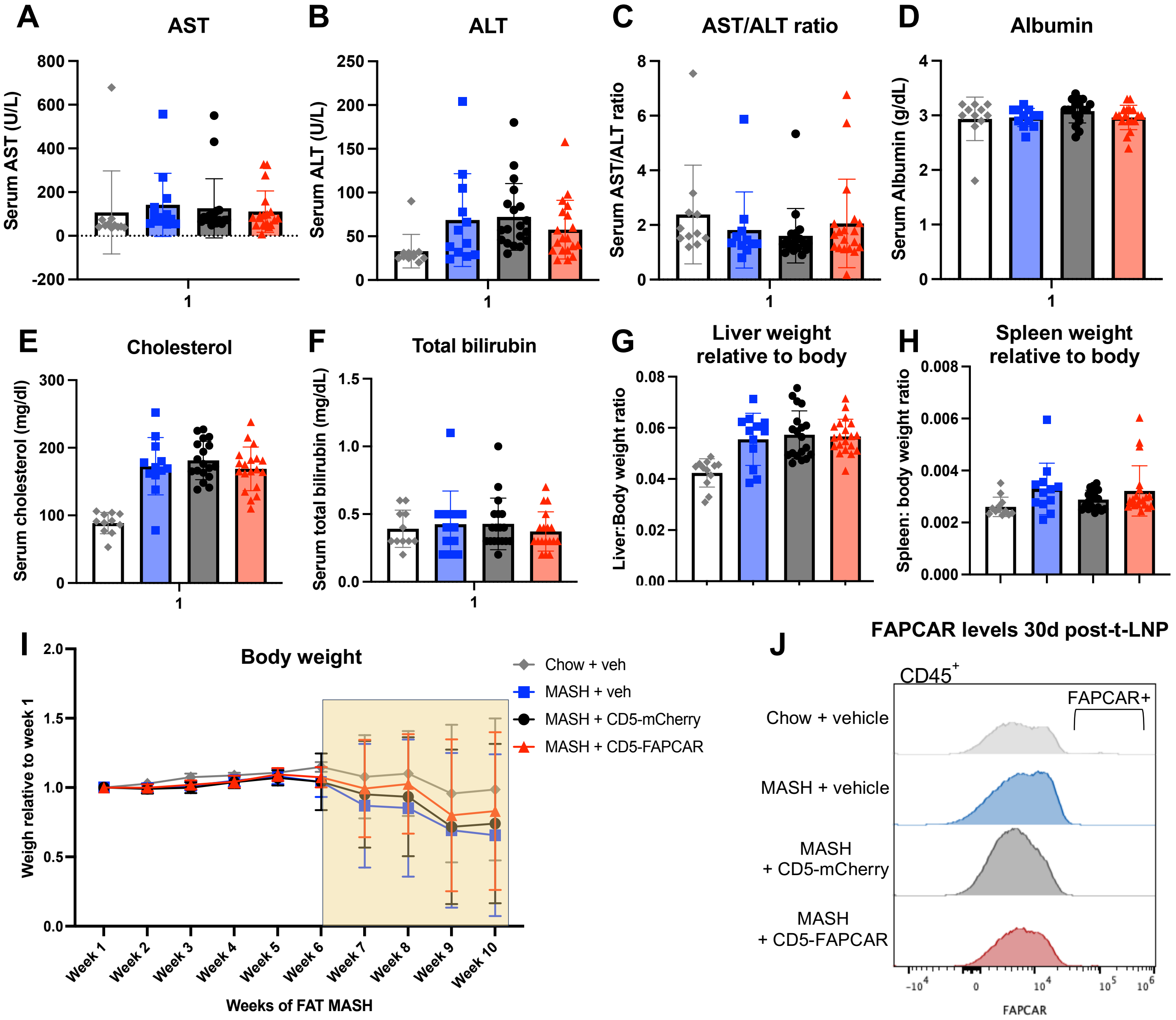


**Fig. S6. One month after t-LNP injection, in vivo anti-FAP CAR T cells are well tolerated and undetectable. (A-H)** Serum values. **(I)** Mouse body weight relative to week 1, where yellow box indicates weeks continued on FAT MASH model post-t-LNP injection. **(J)** Flow plot of FAPCAR levels in CD45+ cells 30 days after t-LNP injection.
