## Supplemental Figure 7 for "In vivo anti-FAP CAR T therapy reduces fibrosis and restores liver homeostasis in metabolic dysfunction-associated steatohepatitis"

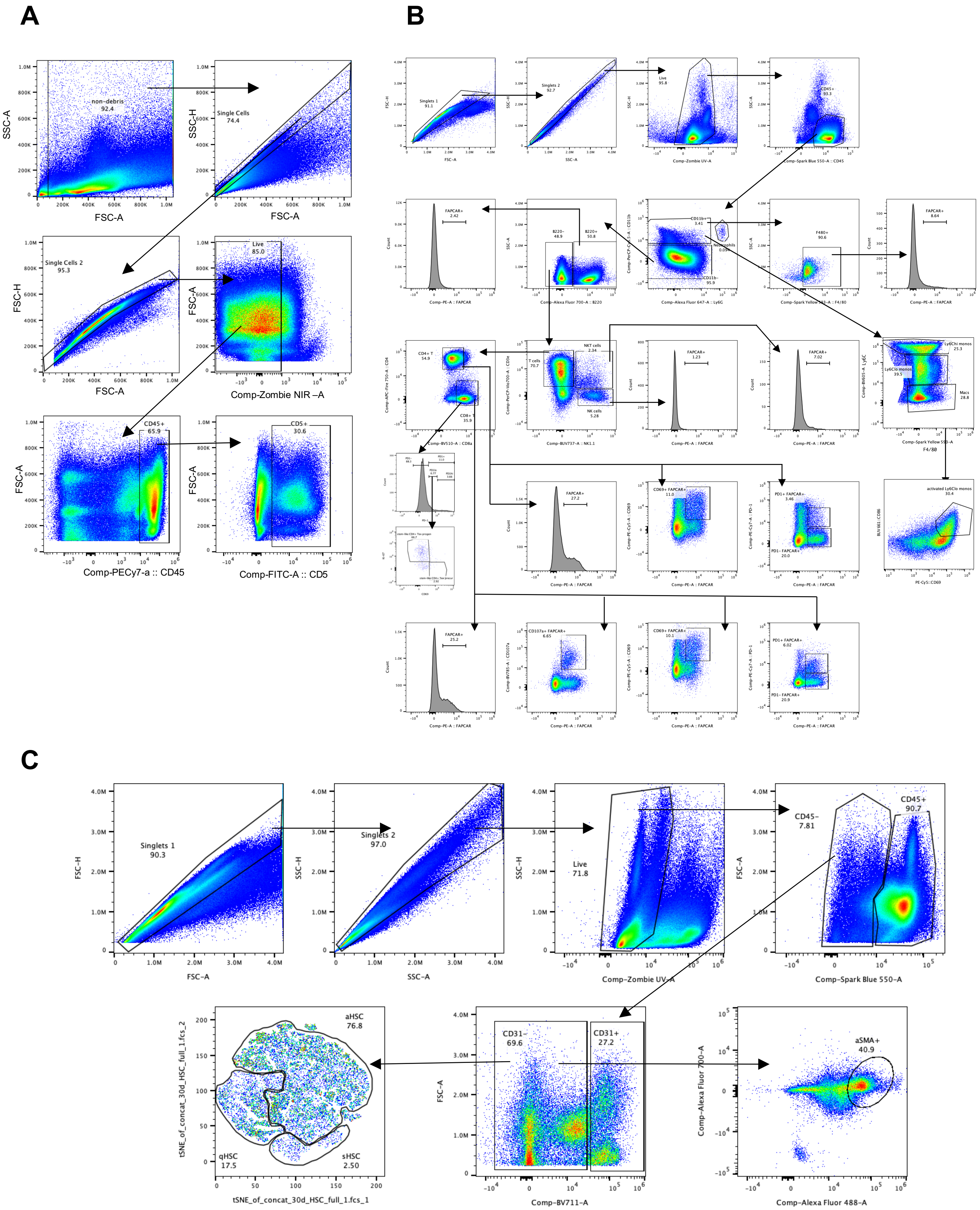


**Fig. S7. Flow gating strategies. (A)** FAPCAR$^+$ immune cell gating. **(B)** Extracellular CD5 gating. **(C)** HSC gating for tSNE plots and αSMA+ HSCs.
