## Supplemental Figure 8 for "In vivo anti-FAP CAR T therapy reduces fibrosis and restores liver homeostasis in metabolic dysfunction-associated steatohepatitis"

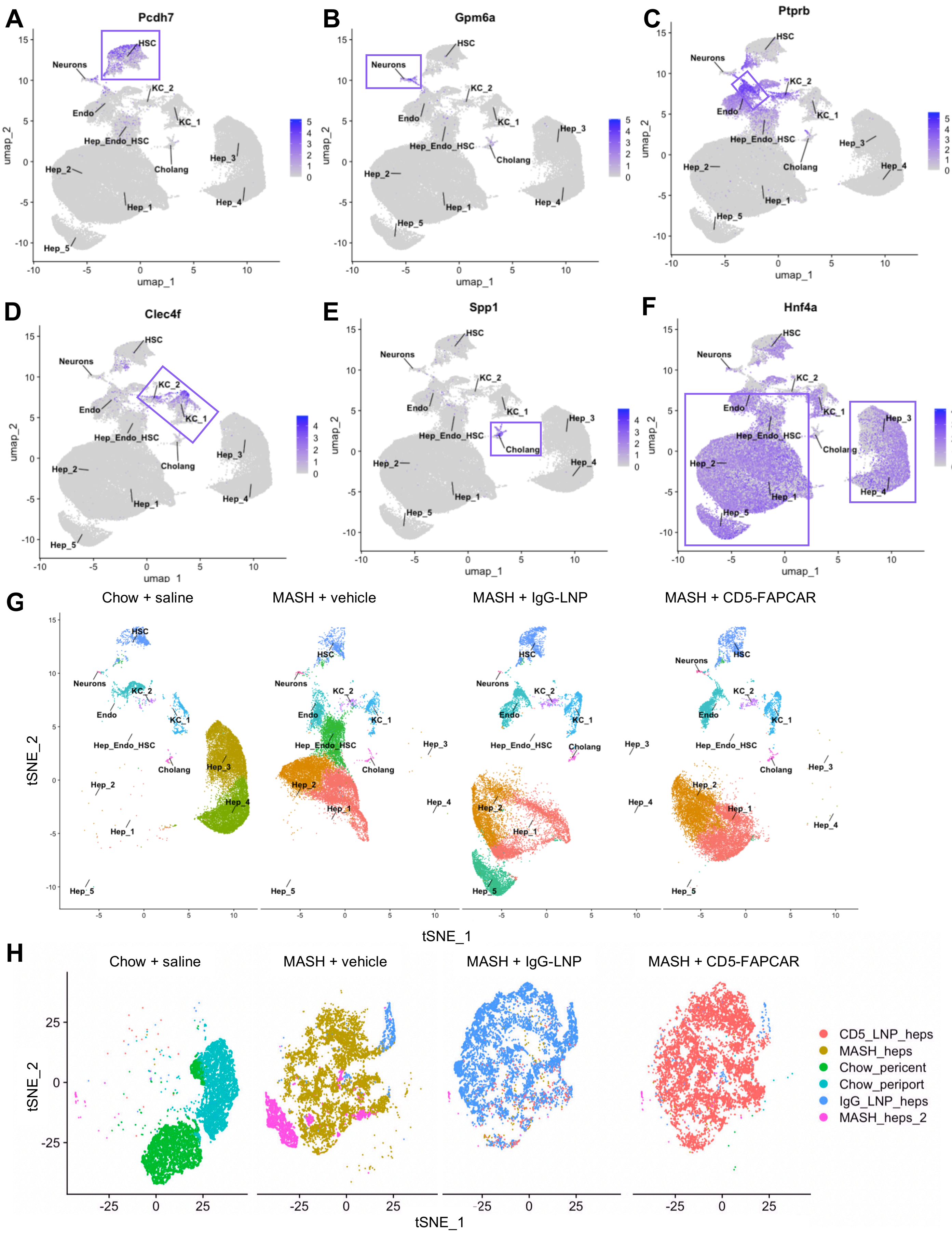

A
Pcdh7
B
Gpm6a
C
Ptprb
D
Clec4f
E
Spp1
F
Hnf4a
HSC
Neurons
KC_2
KC_1
Endo
Hep_Endo_HSC
Cholang
Hep_1
Hep_2
Hep_3
Hep_4
Hep_5
umap_1
umap_2
G
Chow + saline
MASH + vehicle
MASH + IgG-LNP
MASH + CD5-FAPCAR
tSNE_1
tSNE_2
H
Chow + saline
MASH + vehicle
MASH + IgG-LNP
MASH + CD5-FAPCAR
tSNE_1
tSNE_2
CD5_LNP_heps
MASH_heps
Chow_pericent
Chow_periport
IgG_LNP_heps
MASH_heps_2

**Fig. S8. snRNAseq clustering. (A-F)** Example markers used to identify clusters. **(G)** Clusters split by treatment group. **(H)** Hepatocyte populations split by treatment group. All snRNAseq data are from Chow + saline, MASH + saline, MASH + IgG-FAPCAR and MASH + CD5-FAPCAR experiment.
