## Supplemental Table 1 for "In vivo anti-FAP CAR T therapy reduces fibrosis and restores liver homeostasis in metabolic dysfunction-associated steatohepatitis"

**Table S1. Flow cytometry antibodies**

| Antibody | Product # |
|---|---|
| Zombie UV | Biolegend 423107 |
| NK1.1 BUV 737 | BD 741715 |
| Ki67 BV421 | Biolegend 652411 |
| FoxP3 PacBlue | Biolegend 126409 |
| CD8a BV510 | Biolegend 100751 |
| Ly6C BV605 | Biolegend 128035 |
| CD25 BV711 | Biolegend 102409 |
| CD11c BV785 | Biolegend 117335 |
| CD45 Spark Blue 550 | Biolegend 103165 |
| CD11b PerCP/Cy 5.5 | Biolegend 101227 |
| CD3e PerCP-Vio700 | Miltenyi 130-123-282 |
| CD172a RB780 | BD 755521 |
| Ly6G AF647 | Biolegend 127609 |
| B220 AF700 | Biolegend 103232 |
| CD4 APC/Fire 750 | Biolegend 100459 |
| F4/80 Spark YG 593 | Biolegend 157311 |
| CD206 PE/Dazzle 594 | Biolegend 141731 |
| CD69 PeCy5 | Biolegend 104509 |
| CD86 BUV661 | BD Bioscience 741528 |
| PD-1 PE/Cy7 | Invitrogen 25-9985-80 |
| TIM-3 PE/Fire 810 | Biolegend 119745 |
| CD107a BV785 | Biolegend 121641 |
| aSMA AF488 | invitrogen 53-9760-82 |
| CD38 PE/Fire 640 | Biolegend 102743 |
| CD31 BV711 | Biolegend 102449 |
